# Alcohol Use Disorder Associated Gene *FNDC4* Alters Glutamatergic and GABAergic Neurogenesis

**DOI:** 10.1101/2025.05.15.654319

**Authors:** Xiujuan Zhu, August J. John, Li Wang, Sooan Kim, Enci Ding, Ateka Saleh, Irene Marín-Goñi, Abedalrahman Jomaa, Huanyao Gao, Ching Man Wai, Irene Moon, Brandon J. Coombes, Tony M. Kerr, Nobuyoshi Suto, Liewei Wang, Mark A. Frye, Joanna M. Biernacka, Victor M. Karpyak, Hu Li, Richard M. Weinshilboum, Duan Liu

**Author notes:** Corresponding Authors: Duan Liu, Department of Molecular Pharmacology & Experimental Therapeutics Mayo Clinic, 200 First Street SW, Rochester, MN 55905, Richard M. Weinshilboum, Department of Molecular Pharmacology & Experimental Therapeutics Mayo Clinic, 200 First Street SW, Rochester, MN 55905. These authors contributed equally to the study (Xiujuan Zhu, August J. John).

## Abstract

Large-cohort genome-wide association studies (GWAS) for alcohol use disorder (AUD) and AUD-related phenotypes have identified more than one hundred genetic loci. Functional study of those GWAS-identified loci might represent an important step toward understanding AUD pathophysiology. We found that genetic loci which are splicing quantitative trait loci (sQTLs) for the *fibronectin III domain containing 4* (*FNDC4*) gene in the brain were identified by GWAS for both AUD drug treatment outcomes and AUD risk. However, FNDC4 function in the brain and how it might contribute to AUD pathophysiology remain unknown. In the present study, we characterized GWAS locus-associated *FNDC4* splice isoforms, studies which suggested that FNDC4 alternative splicing results in loss-of-function for FNDC4. We also investigated *FNDC4* function using CRISPR/cas9 gene editing, and the creation of human induced pluripotent stem cell (iPSC)-derived neural organoids joined with single-nucleus RNA sequencing. We observed that knock-out (KO) of *FNDC4* resulted in a striking shift in the relative proportions of glutamatergic and GABAergic neurons in iPSC-derived neural organoids, suggesting a possible important role for FNDC4 in neurogenesis. We also explored potential mechanism(s) of *FNDC4*-dependent neurogenesis with results that suggested a role for FNDC4 in mediating neural cell-cell interaction. In summary, this series of experiments indicates that FNDC4 plays a role in regulating cerebral cortical neurogenesis in the brain. This regulation may contribute to the response to AUD pharmacotherapy as well as the effects of alcohol on the brain.

## INTRODUCTION

Alcohol use disorder (AUD) is the most common addictive disorder and a leading cause of disease burden worldwide^1, 2^. Patients with AUD experience intense alcohol craving that drives increased alcohol ingestion despite harmful health consequences^1, 2^. It is believed that AUD may develop as a result of disrupted homeostasis of neurotransmission after exposure to alcohol which can influence multiple brain neurotransmitter systems^3^. However, not every subject who drinks alcohol will develop AUD, indicating that susceptibility to this disorder is multifactorial. Genetics clearly contributes to AUD risk^4, 5^. Recently large-cohort human genomic studies of AUD and AUD-related phenotypes have identified a large number of genetic loci^6–11^, results which have significantly advanced our understanding of the genetic etiology of AUD. Some of those GWAS-identified loci appeared to be aligned with known mechanisms of AUD pathophysiology, *e.g.*, alcohol metabolism (*ADH1B, ADH1C*) and neurotransmission (*DRD2, GABRA4, OPRM1, CACNA1C*). However, like many GWAS, the majority of AUD GWAS-identified loci mapped to non-coding regions, and their function in AUD needs to be investigated further^5^.

Only three medications are currently approved by the U.S. Food and Drug Administration for use in the treatment of AUD, with acamprosate and naltrexone being far and away the most widely prescribed^2, 12^. Both drugs are believed to target mechanisms of neurotransmission. However, response to treatment with these medications is variable, with only ∼50% of AUD patients achieving significant positive treatment outcomes^12, 13^. In an effort to better understand the contribution of genetic factors to individual variation in response to AUD pharmacotherapy, we performed a genome-wide association study (GWAS) for treatment outcomes that included over one thousand AUD patients who received acamprosate and/or naltrexone therapy^13^. That GWAS identified a genome-wide significant single-nucleotide polymorphism (SNP) locus that colocalized with a splicing quantitative trait locus (sQTL) for the *fibronectin III domain containing 4* (*FNDC4*) gene in multiple human brain regions^14^. We also noted an independent SNP locus which is also an sQTL for *FNDC4* in the human brain^14^ and which has been repeatedly associated with AUD risk and alcohol consumption in large-cohort GWAS^6–11^. The fact that brain *FNDC4* sQTL SNPs are significantly associated with both AUD risk and AUD drug treatment outcomes strongly suggests a possible role for brain FNDC4 in AUD pathophysiology. However, in spite of the fact that *FNDC4* is highly expressed in the brain^14^, its function in the central nervous system (CNS) and how that function might contribute to AUD pathophysiology remain unknown.

In the present study, we set out to characterize the molecular function of *FNDC4* in the CNS with the goal of understanding its possible role in AUD pathophysiology and its contribution to variation in AUD drug treatment response. We began with functional annotation and characterization of the GWAS SNP-associated *FNDC4* RNA splice isoforms, studies which suggested that the GWAS SNPs might be associated with FNDC4 decrease or loss-of-function in the brain. As the next step, we used CRISPR/cas9 gene editing, the creation of human induced pluripotent stem cell (iPSC)-derived dorsal forebrain organoids, and single-nucleus RNA sequencing (snRNA-seq) to show that *FNDC4* knock-out increases glutaminergic neurons and decreases GABAergic neurons in the organoids. This series of observations suggests that FNDC4 may play a role in regulating glutamatergic and GABAergic neurogenesis—at least at certain early stages of neurodevelopment, a function which may contribute to the effect of alcohol on the brain as well as clinical response to alcohol exposure and AUD drug therapy.

## RESULTS

### Potential association of brain FNDC4 alternative splicing with AUD phenotypes

Our previous GWAS for AUD drug treatment outcomes identified a genome-wide significant SNP signal (“top” SNP, rs56951679, T>C; *p*=1.6×10^−8^) that was associated with “time until heavy relapse to alcohol drinking” after 90 days of acamprosate and/or naltrexone therapy^13^. The minor allele “C” was associated with worse treatment outcomes (**Fig. 1A**). The rs56951679 SNP locus colocalized with an *FNDC4* sQTL in multiple brain regions including cortex, nucleus accumbens and three other brain regions based on GTEx data^14^ (**Fig. 1B, Supplemental Fig. S1**). The minor allele, “C”, was associated with increased intron/exon ratio at splice sites (*chr2*:27,493,389-27,472,478bp) that excise exon 6 of the FNDC4 reference mRNA (**Fig. 1B, 1C**), indicating an increased level of the FNDC4 variant mRNA splice isoform in the brains of individuals who carry the “C” allele. In addition, large-cohort GWAS for AUD risk and alcohol consumption have repeatedly identified a different SNP locus (index SNP, rs1260326, C>T)^6–11^ which also colocalized with an sQTL for *FNDC4* in the human brain^14^ (**Supplemental Fig. S2**). That rs1260326 SNP was not linked to the rs56951679 SNP in European populations (*r*^2^ < 0.033, D’ <0.445, based on 1000 Genomes data). The rs1260326 SNP common allele “C” that was associated with increased AUD risk in large-cohort GWAS^6–11^, was also associated with increased intron/exon ratio at splice sites which exon 6 had been deleted (**Supplemental Fig. S2**).

**Figure 1.**
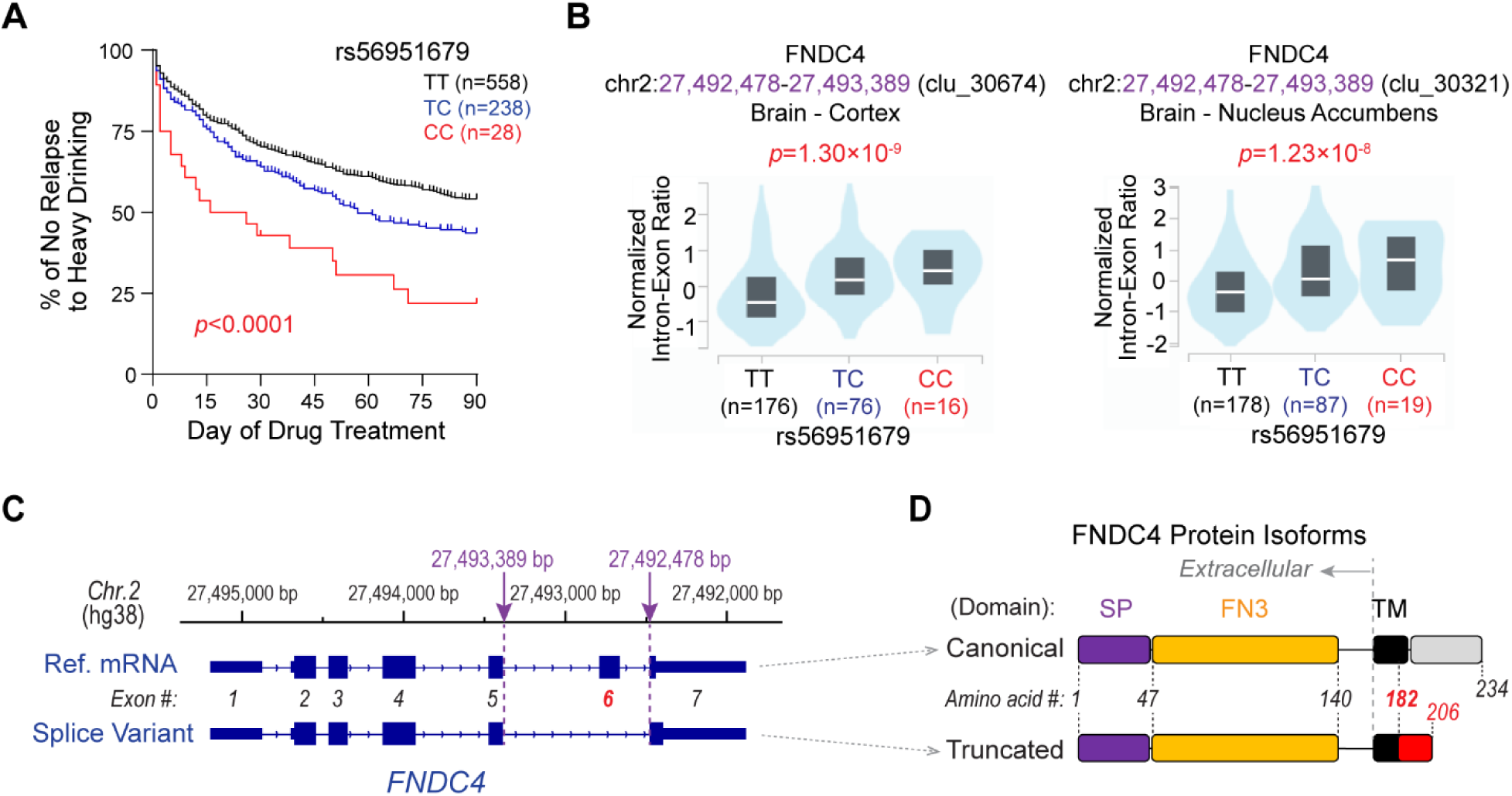
AUD drug treatment response and *FNDC4* alternative splicing in the brain. (**A**) Kaplan-Meier plot showing the percentage of AUD patients who did not relapse to heavy alcohol drinking during 90 days of pharmacotherapy. A smaller percentage of patients who carried the rs56951679 SNP variant allele “C” remained abstinent from heavy alcohol drinking, suggesting that worse drug treatment response is seen in patients carrying the rs56951679 SNP “C” allele. The *p*-value was calculated using Log-rank (Mantel-Cox) test. KM plot was generated based on our published GWAS data^13^. (**B**) The rs56951679 SNP is an sQTL for the *FNDC4* gene in many different human brain regions. The SNP genotype was associated with FNDC4 RNA splicing in brain cortex (left) and nucleus accumbens (right) based on the RNA-seq data generated by GTEx (v10)^14^. The variant allele “C” was associated with an increased level of intron excision on chromosome 2 (*chr.2*): 27,492,478-27,493,389 (hg38). (**C**) Depiction of the rs56951679 SNP-associated alternative splice sites and FNDC4 RNA splice isoforms. Panels from top to bottom are (top panel) physical position of the *FNDC4* gene on *chr.2* based on human genome assembly hg38 with sQTL splice sites highlighted in purple; The reference (Ref.) FNDC4 mRNA with 7 exons is numbered below (middle panel); as is the FNDC4 splice variant with exon 6 excision (bottom panel). FNDC4 transcripts were annotated by the RefSeq. (**D**) Depiction of the FNDC4 protein encoded by the Ref. (canonical) and spliced (truncated) FNDC4 RNA isoforms. The canonical FNDC4 protein includes a signal peptide (SP), a fibronectin type III domain (FN3), a transmembrane domain (TM) and a cytosolic C-terminal. The truncated FNDC4 protein displays a frameshift after amino acid 182 which disrupts the TM domain.

Both our GWAS for AUD drug treatment response^13^ and the GWAS for AUD risk^6–11^ identified SNP loci that colocalized with *FNDC4* sQTLs in brain tissue, results which suggested a potential correlation between brain FNDC4 splicing and AUD clinical phenotypes. Specifically, increased levels of the variant FNDC4 splice isoform in the brain were associated with worse AUD drug treatment outcomes as well as elevated AUD risk. In addition, a previous transcriptome-wide study reported that *Fndc4* was one of the top 6 genes for which mRNA expression levels were significantly decreased in the mouse brain during alcohol withdrawal^15^. Taken together, these human and mouse multi-omics data strongly support a possible role for variation in brain-expression of FNDC4 in alcohol-related phenotypes. However, in spite of the fact that *FNDC4* is highly expressed in the brain^14^, most highly in neurons when compared to other cell types^16^, its function in the CNS remains unknown. Therefore, we set out to characterize *FNDC4* function to better understand biological mechanism(s) that might contribute to its role in AUD-related phenotypes.

### FNDC4 splice isoforms encode a truncated protein with the loss of cell membrane localization and of glycosylation

Our functional studies began with characterization of the GWAS-SNP-associated FNDC4 splice isoform with the loss of exon 6 when compared to the reference mRNA (**Fig. 1C**). As encoded by reference mRNA, the canonical FNDC4 protein included a signal peptide (SP), a fibronectin type III domain (FN3), a transmembrane domain (TM) and a cytosolic C-terminal domain (see **Fig. 1D**). As a result, it was predicted to be a single-pass type I transmembrane protein with three putative *N*-linked glycosylation sites according to UniProt^17^. The loss of exon 6 in the FNDC4 mRNA splice isoform resulted in a truncated protein, a change in encoded amino acid sequence that interrupted part of the TM domain (see **Fig. 1D**). This change could potentially affect FNDC4 cell membrane localization.

As a first step designed to determine whether the truncated protein might or might not translocate to the cell membrane, we overexpressed both the canonical and truncated FNDC4 proteins in HEK293 cells with MYC- and FLAG-tags fused to their C-termini (**Fig. 2A**). Molecular weights (MW) for the overexpressed MYC-FLAG fused FNDC4 proteins were approximately 3 kD larger than the predicted MWs for the native proteins which were 25.2 kD (canonical) and 22.7 kD (truncated) (**Fig. 2A**). We validated overexpression by Western blot analysis using either FLAG (**Fig. 2B**, left panel) or MYC antibodies (**Supplemental Fig. S3A**). We observed multiple bands for the overexpressed canonical FNDC4 protein. One of those bands matched its predicted MW of 28 kD (**Fig. 2B**, black arrow) while other bands had higher MWs close to 37 kD (**Fig. 2B**, blue arrow). By contrast, only one major band was observed for the truncated protein, a band with a MW that matched its predicted size (**Fig. 2B**, red arrow). Using those same overexpressed canonical and truncated FNDC4 protein samples as positive controls, we also tested multiple commercially available anti-FNDC4 antibodies (**Supplemental Fig. S3B, 3C**; and **Supplemental Table S1**). The most efficient anti-FNDC4 antibody (Ab-4, see **Fig. 2B**, right panel) was then used to study endogenous FNDC4 protein in brain tissue.

**Figure 2.**
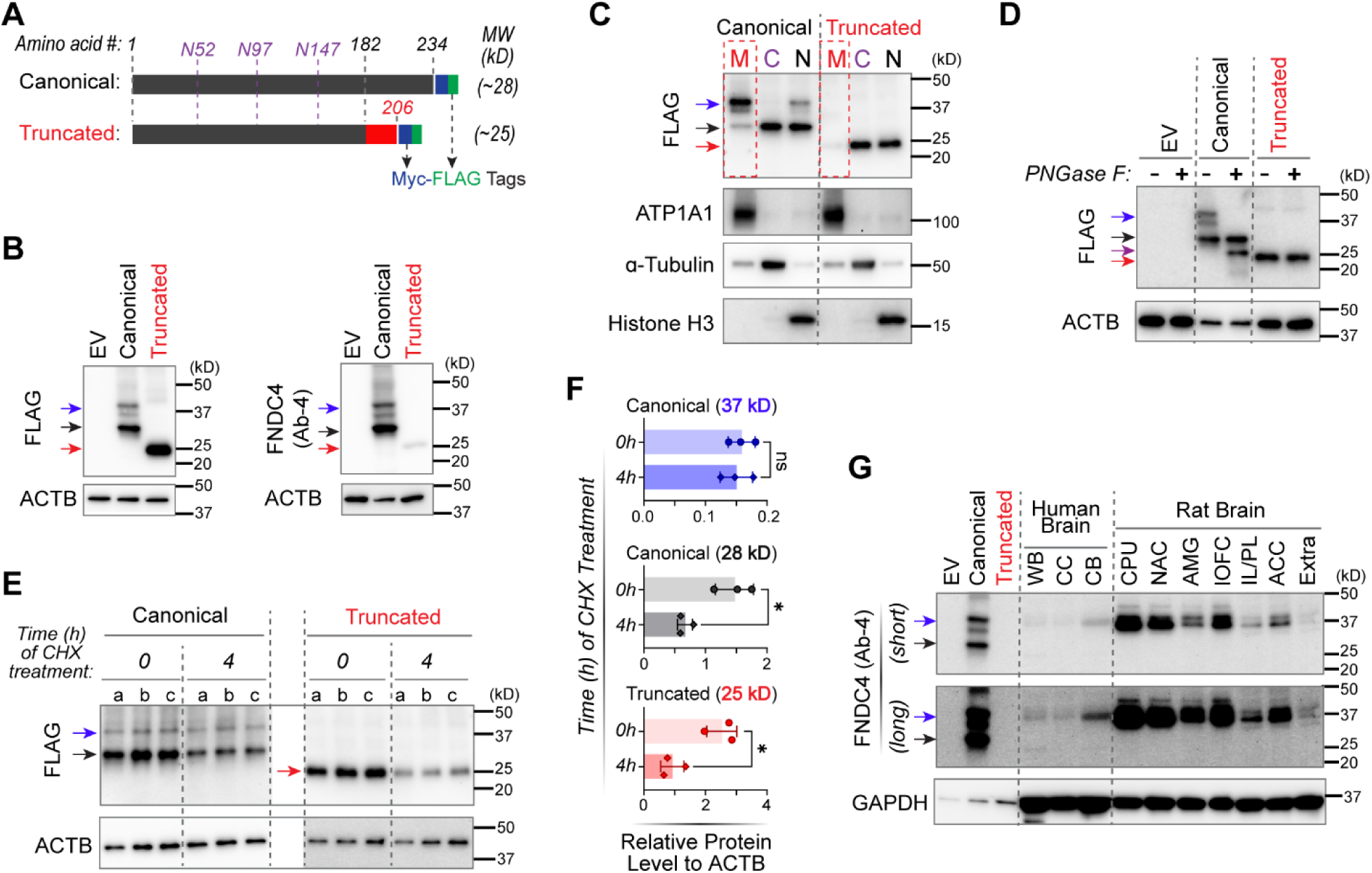
FNDC4 splice isoform characterization. (**A**) Depiction of the cDNA-overexpressed canonical and truncated FNDC4 proteins with MYC- and FLAG-tags fused at their C-termini. The number of amino acids was labelled on the top panel. Three asparagine (N) which are putative *N*-linked glycosylation sites were labelled in purple. The predicted molecular weights (MWs) of both overexpressed FNDC4 proteins were also listed. (**B**) The Western blot shows cDNA-overexpressed canonical and truncated FNDC4 proteins in HEK293T cells. Anti-FLAG antibody (left panel) was first tested to ensure the overexpression of both protein isoforms. Compared with the empty vector (EV) control, canonical FNDC4 protein showed multiple bands (blue & black arrows) while the truncated protein showed only one major band (red arrow, top panel). Those same protein samples were then used to test anti-FNDC4 antibodies. One of those tested FNDC4 antibodies (Ab-4) and showed the best efficiency and specificity in detecting the overexpressed FNDC4 proteins (right panel). Beta-actin (ACTB) was blotted as a loading control. (**C**) Blots for overexpressed FNDC4 protein isoforms in subcellular fractions of the membrane (M), cytoplasm (C) and nucleus (N). The larger bands (blue arrow) of canonical FNDC4 map mainly to the membrane while the smaller band (black arrow) maps mainly to the cytoplasm and nucleus. Truncated FNDC4 (red arrow) was located mainly in the cytoplasm and nucleus but not in the membrane. ATP1A1, α-tubulin, and histone H3 were blotted as markers for the cell membrane, cytoplasm and nucleus, respectively. (**D**) FNDC protein deglycosylation assay visualized by Western blots. Overexpressed FNDC4 protein isoforms were treated with the peptide-*N*-glycosidase F (PNGaseF) that cleaves oligosaccharides from *N*-linked glycoproteins. Compared to the non-treatment control (-), PNGaseF treatment (+) removed the larger bands (blue arrow) of canonical FNDC4, and simultaneously a smaller band emerged (purple arrow). The truncated FNDC4 band was not affected by PNGaseF treatment. (**E**) Western blots for overexpressed FNDC4 proteins in HEK293 cells after 0- and 4-hours of treatment with cycloheximide (CHX) which inhibits protein synthesis. Lanes a, b and c are biological triplicates. (**F**) Relative protein levels of overexpressed FNDC4 to ACTB. Protein level was quantified based on band intense showed in (E). Each dot or diamond represents one of biological triplicate samples with 0- or 4-hours CHX treatment, respectively. Significant decreases in un-glycosylated canonical (28 kD) and truncated FNDC4 proteins (25 kD) were observed but not in glycosylated FNDC4 (37 kD) after 4-hours CHX treatment. \**p* <0.05 and ns =not significant by Student’s *t* test. (**G**) Western blots for FNDC4 in human and rat brain lysates. Overexpressed canonical and truncated FNDC4 proteins were blotted as positive controls. Protein lysates of human whole brain (WB), cerebral cortex (CC) and cerebellum (CB) were obtained from a commercial source (see Table S1). Protein lysates of different rat brain regions included caudate putamen (CPU), nucleus accumbens (NAC), amygdala (AMG), lateral orbitofrontal cortex (lOFC), infralimbic/prelimbic cortices (IL/PL), anterior cingulate cortex (ACC), and “extra” forebrain tissues containing medial orbitofrontal cortex and motor cortex (Extra) were prepared from snap frozen fresh tissues of three rats. GAPDH was blotted as an internal control.

We next determined the subcellular localization of both the canonical and truncated FNDC4 proteins by Western blot analysis after subcellular fractionation of cell membrane, cytoplasm and nucleus of HEK293T cells that overexpressed FNDC4 protein. For the canonical FNDC4 protein, we found that the ∼37 kD bands (**Fig. 2C**, blue arrow) were predominantly present in the cell membrane (M) and that the ∼28 kD band (**Fig. 2C**, black arrow) was present in the cytoplasm (C) and nucleus (N). By contrast, the truncated FNDC4 protein was present mainly in the cytoplasm and nucleus but **not** in the cell membrane (**Fig. 2C**, red arrow). These results demonstrated that the canonical FNDC4 protein could translocate to the cell membrane but that the truncated protein, encoded by the GWAS-SNP-associated FNDC4 splice variant (**Fig. 1C, D**), could not efficiently translocate to the cell membrane.

FNDC4 has been studied in peripheral tissues but the Western blot results for those studies are contradictory, with two studies reporting multiple bands^18, 19^ and one reporting only a single band^20^. Since we had observed multiple bands for the canonical FNDC4 protein (**Fig. 2B**) and, most importantly, since those bands were distributed at different subcellular localizations (**Fig, 2C**), we next investigated what those multiple bands might represent. As demonstrated subsequently, these studies were critical for our understanding of the function of both canonical and truncated FNDC4 proteins. We found that the cell membrane-enriched bands (**Fig. 2D**, blue arrow) represented glycosylated canonical FNDC4 protein because those bands disappeared, and a smaller band appeared (**Fig. 2D**, purple arrow), after the protein sample had been treated with PNGaseF, an endoglycosidase which can cleave *N*-linked oligosaccharides from glycoproteins^21^. By contrast, the truncated FNDC4 protein band was not affected by PNGaseF exposure (**Fig. 2D**, red arrow), suggesting that the truncated protein was not glycosylated as was the canonical protein. This result was consistent with bioinformatic predictions that FNDC4 had three putative *N*-linked glycosylation sites (**Fig. 2A**), and the fact that glycoproteins often show higher MWs in Western blot than their predicted MWs^22–24^. The MW of the newly-appearing band (**Fig. 2D**, purple arrow) was smaller than the cytoplasm-enriched band (**Fig. 2D**, black arrow), which likely was due to both de-glycosylation by endoglycosidases and cleavage of the signal peptide (SP) (**Fig. 1D**) since the ∼37 kD bands (blue arrow) were enriched in the membrane and since SP cleavage often occurs when a newly synthesized protein is translocated to the membrane^25^.

We also observed that the truncated and the un-glycosylated canonical FNDC4 proteins (**Fig. 2E**, red & black arrows) both of which were not glycosylated, were less stable than was glycosylated FNDC4 (**Fig. 2E**, blue arrow). Specifically, when HEK293 cells overexpressing FNDC4 protein were treated with cycloheximide (which inhibits protein synthesis) for 4 hours, significant decreases in un-glycosylated canonical and truncated FNDC4 proteins were observed, but that was not the case for glycosylated FNDC4 (**Fig. 2E, 2F**).

In summary, this series of experiments demonstrated that the GWAS-SNP-associated FNDC4 splice variant isoform encoded a truncated protein which could not be efficiently glycosylated or translocated to the cell membrane, resulting in an unstable and rapidly degraded protein in the cytoplasm when compared with the glycosylated canonical protein. This result suggested that the GWAS-SNP-associated FNDC4 splice variant in brain might have contributed to AUD-related phenotypes because it encoded an unstable truncated protein which compromised FNDC4 function in the brain. This series of experiments had characterized FNDC4 protein isoforms in detail, a crucial initial step for the study of endogenous FNDC4 in the brain, as detailed subsequently.

### FNDC4 is primarily a transmembrane glycoprotein in the brain

Before studying FNDC4 function using CNS models, we attempted to detect endogenous FNDC4 protein isoforms in the human brain since that information would help us understand its function in the brain. We used a validated FNDC4 antibody (**Fig. 2B**) to detect FNDC4 proteins in commercially available human brain protein lysates by Western blot. Those experiments showed major band(s) with MW which matched that of the glycosylated canonical FNDC4 protein present in human whole brain (WB), cerebral cortex (CC) and cerebellum (CB) lysates (**Supplemental Fig. S3D**).

To further characterize endogenous FNDC4 protein isoforms in brain, we harvested both cortical and subcortical regions of rat brains and snap froze them to obtain protein lysates. This step made protein lysates from fresh brain tissue available which we could not obtain from human subjects. FNDC4 protein sequences are highly conserved across mammalian species. For example, human and rat FNDC4 protein sequences are almost identical with less than 5% differences which are found in the signal peptide (see **Supplemental Fig. S4**), a domain which is often cleaved after proteins are translocated to the cell membrane^25^. Therefore, we anticipated that the human FNDC4 antibody could also detect rat Fndc4 during Western blot analysis. In fact, we detected endogenous Fndc4 bands in multiple rat brain regions (**Fig. 2G**). Those Fndc4 bands were more intense than those detected in human brain lysates, observations which might be due to the method used to prepare and store protein lysates. Of importance, the MWs of FNDC4 bands in both human and rat brain lysates were similar (∼37 kD) and they matched well with the glycosylated membrane-enriched canonical FNDC4 protein (**Fig. 2G**, blue arrow). By contrast, the un-glycosylated cytoplasm-enriched canonical FNDC4 protein (**Fig. 2G**, black arrow) was barely detectable in either human or rat brain lysates.

Of note, although this FNDC4 antibody (Ab-4) could detect overexpressed truncated FNDC4, its efficiency in detecting truncated FNDC4 was much lower than in detecting canonical FNDC4 (**Fig. 2B**). It was anticipated that this antibody might not be able to detect the truncated FNDC4 if it presents in human and/or rat brain lysates (**Fig. 2G**). However, we have demonstrated that truncated FNDC4 could not be translocated to the cell membrane or glycosylated, resulting in rapid degradation when compared with the canonical glycosylated protein (**Fig. 2C-2F**). Therefore, the truncated FNDC4 protein is unlikely to be functional.

FNDC4 has been reported to be cleaved at the cell membrane with release of its extracellular portion as a soluble peptide^18^. That conclusion was based on cDNA overexpression experiments similar to those performed here (**Fig. 2A**). Those authors overexpressed a FLAG-tag (N-terminus) and MYC-tag (C-terminus) fused mouse Fndc4 protein with an amino acid sequence which is very similar to that of human FNDC4 (see **Supplemental Fig. S4**). They also detected multiple bands of the overexpressed Fndc4 protein by Western blot analysis of cell lysates, just as we did for our human FNDC4 canonical protein (**Fig. 2B**). In addition, they observed only a single band (∼25kD) in the cell culture media using antibody directed against an N-terminus fused FLAG-tag^18^. Those investigators concluded that the band detected in the media (∼25kD) represented a “cleaved and secreted” portion of their overexpressed Fndc4 protein because “the band was of reduced MW compared with the cellular version” (although without cell lysates as a control in the same blot) and was undetectable in the media when they tested antibody against a C-terminus fused MYC-tag^18^. However, those results do not exclude the possibility that the single band detected in cell culture media was an un-glycosylated cytoplasm-enriched full-length Fndc4 which could be released into cell culture media, as we observed in a similar experiment (**Supplemental Fig. S3E**).

Taken as a whole, our results strongly suggest that the functional endogenous FNDC4 protein in brain is primarily a transmembrane glycoprotein. That information is important for the experiments designed to investigate FNDC4 function in brain, as described subsequently.

### *FNDC4* knock-out alters glutamatergic and GABAergic neurogenesis and their relative balance in human iPSC-derived neural organoids

The function of FNDC4 in the brain was unclear when we began our studies. However, it is a protein with a simple structure for which the major potential functional domain is an extracellular fibronectin type III (FN3) domain (see **Fig. 1D**). FN3 domains are known to bind to protein partners and to mediate cell-cell interaction^26^. In addition, we have demonstrated that FNDC4 in brain is primarily a transmembrane glycoprotein (**Fig. 2**). Those facts led us to the hypothesis that FNDC4 might function through cell surface protein binding and the mediation of neural cell-cell interaction. Therefore, as the next step in our studies, we addressed the issue of human FNDC4 function using induced pluripotent stem cell (iPSC)-derived neural organoids, which might provide an opportunity to capture and study FNDC4 function because those organoids are self-assembled 3-dimensional structures containing multiple neural cell types and they can partially capture cell-cell interactions during processes of human brain development^27–29^. Specifically, we generated *FNDC4* knock-out (KO) iPSCs by CRISPR/cas9 and differentiated both wild-type (WT) and homozygous *FNDC4* KO iPSCs to generate dorsal forebrain organoids. Those organoids were collected and studied at different timepoints (45, 90 and 150 days) of differentiation/maturation and were also analyzed by snRNA-seq (see **Fig 3A**).

**Figure 3.**
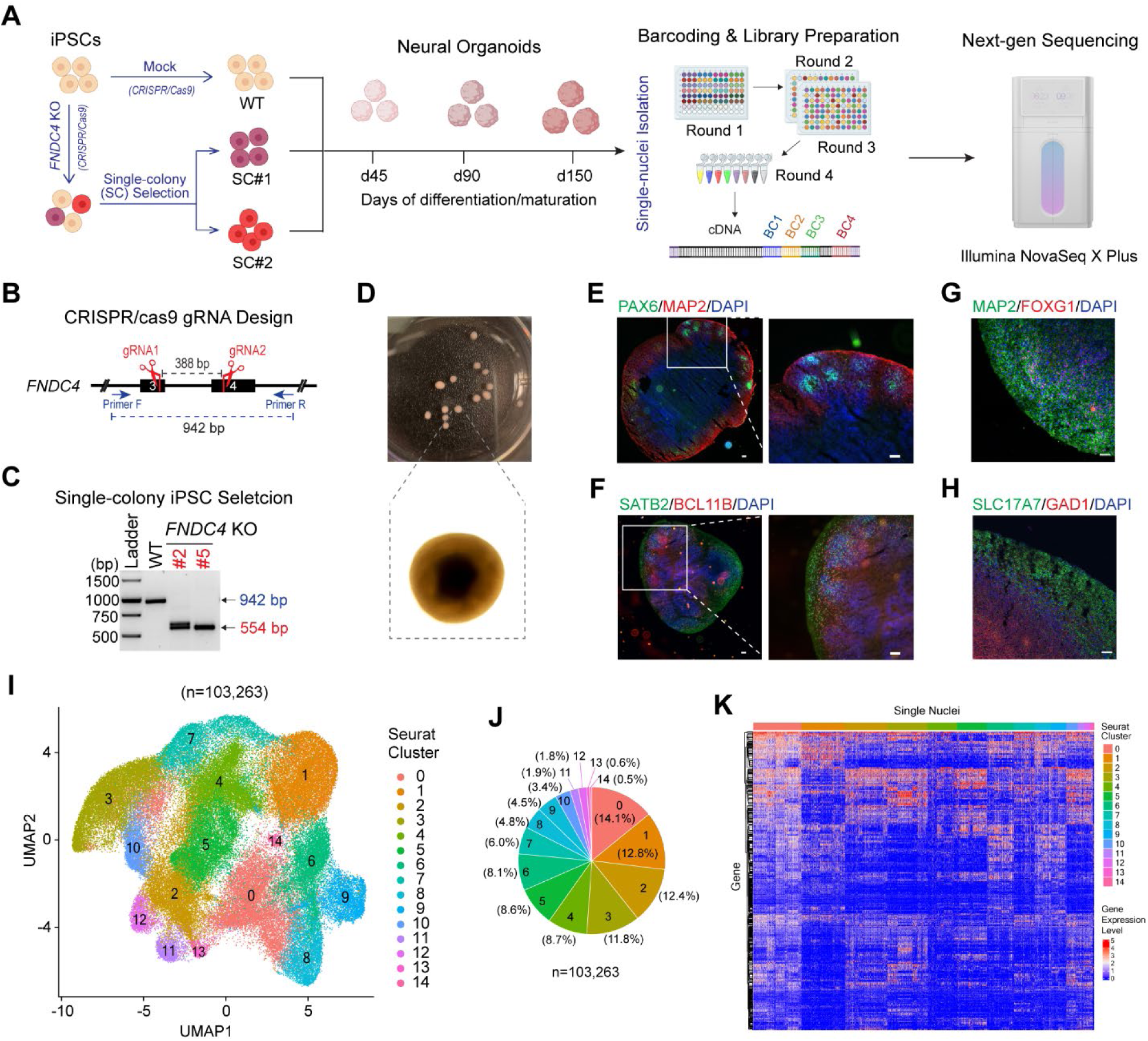
FNDC4 functional study using human iPSC-derived forebrain organoids. (**A**) Depiction of experimental design. A human iPSC line was used to knock-out (KO) the *FNDC4* gene by CRISPR/cas9. Mock KO was performed without using guide RNAs (gRNAs) which served as a wildtype (WT) control. Two homozygous KO single-colony (SC) iPSC lines were selected and differentiated to forebrain organoids as well as the WT iPSC. Three organoids generated from each iPSC line were harvested at three time points (day 45, 90 and 150). A total of 27 organoids (3 organoids × 3 iPSC lines × 3 time points) were used for single-nucleus isolation. RNA in those single-nuclei was barcoded using “split-pool” combinatorial barcoding technology and were reverse transcribed to cDNA for sequencing using the Illumina NovaSeq X plus platform. (**B**) Design of gRNAs for *FNDC4* CRISPR/cas9 editing, and of PCR primers for *FNDC4* KO colony selection. Successful CRISPR/cas9 editing using both gRNAs were expected to remove 388 base pairs (bp) from the *FNDC4* gene, resulting in a 554 bp amplicon by PCR using designed primers. (**C**) Agarose gel results showing the PCR amplicons using genomic DNA from the WT and two selected *FNDC4* KO iPSC lines. See *Supplemental Fig. S5* for details with regard to iPSC single-colony selection, genotyping and karyotyping. (**D**) iPSC-derived forebrain organoids in one well of a 6-well plate at day 90 of differentiation/maturation (up), and an organoid under the microscope. Representative cryosections of forebrain organoids at day 150 of differentiation/maturation stained for markers of (**E**) neural progenitor cells (PAX6) and neurons (MAP2), (**F**) superficial (SATB2) and deep (BCL11B, also known as CTIP2) neurons, (**G**) developing forebrain neurons (MAP2 and FOXG1), and (**H**) glutamatergic (SLC17A7, also known as VGLUT1) and GABAergic (GAD1, also known as GAD-67) neurons. Scale bars are 100 µM in (E) (F) and 50 µM in (G) (H). See *Supplemental Fig. S6* for details with regard to organoid differentiation/maturation and characterization. (**I**) UMAP visualization of snRNA-seq data for 27 forebrain organoids from 9 experimental conditions. Each dot represents a single nucleus(n=103,263). Nuclei were clustered based on similarity of their transcriptomic profiles (Seurat clustering). (**J**) Pie chart showing proportions of single nuclei in each Seurat cluster. (**K**) Heatmap showing the expression levels of genes that were differentially expressed across different Seurat clusters (color-coded and labelled in the top panel). Each row is a gene, and each column is a single nucleus. For visualization, a total of 5,000 single nuclei were randomly pulled from the single nuclei pool (n=103,263) but with the maintenance of proportions of individual Seurat clusters (J), and the “top” 100 differentially expressed genes in each Seurat cluster based on adjusted *p*-value (Bonferroni correction) were used to plot the heatmap.

To create *FNDC4* KO iPSCs, we designed two guide RNAs (gRNAs) which would remove 388 base pairs (bp) and cause a frameshift in the open reading frame (**Fig 3B**, and **Supplemental Fig. S5**). After CRISPR/cas9 editing using these two gRNAs, selection of single-colony (SC) *FNDC4* KO iPSC lines was performed (see **Supplemental Fig. S5D**). Two of the *FNDC4* KO iPSC lines that maintained the homozygous KO genotype after 3 passages were selected for further study (SC#2 and #5 in **Fig 3C** and in **Supplemental Fig. S5E**) and were characterized by karyotyping to ensure genomic integrity (see **Supplemental Fig. S5F**).

Both *FNDC4* KO iPSC lines as well as WT iPSCs were then differentiated to generate dorsal forebrain organoids based on an established protocol^30, 31^ (see **Supplemental Fig. S6**). We used dorsal forebrain organoids because FNDC4 is most highly expressed in human brain cerebral cortex (GTEx) and because the prefrontal cortex is one of the brain “reward regions” which play important roles in the neurobiology of addiction^32^. The protocol that we used can reliably and reproducibly generate dorsal forebrain organoids with a rich diversity of cell types appropriate for human cerebral cortical tissue^31^. Those iPSCs-derived dorsal forebrain organoids were visually similar in their shape and structure based on immunostaining of neural markers (see **Fig. 3D-H**, and **Supplemental Fig. S6**), all of which were comparable to those used to illustrate the original publications^30, 31^ with regard to these organoids.

To explore their molecular characteristics, the organoids were further analyzed by snRNA-seq. Specifically, three organoids for each experimental condition were pooled for one single-nuclei sample. A total of 9 single-nuclei samples matched with the 9 experimental conditions (3 iPSC lines × 3 time points) were prepared. Nuclei from the 9 samples were barcoded using “split-pool” combinatorial barcoding technology (**Fig. 3A**) which allows sample multiplexing^33^ (Parse Biosciences). Because all 9 samples were barcoded and sequenced in a single experiment, comparisons across samples could be controlled with minimal batch effect. A total of 154,955 single nuclei from 9 samples were sequenced with a mean sequencing depth of ∼20K reads/cell. Single nuclei with fewer than 500 detected genes were filtered out, leaving a total of 103,263 single nuclei for further analysis. The snRNA-seq data were analyzed by Seurat clustering which identified 15 cell clusters (**Fig. 3I**), with proportions ranging from 0.5% to 14.1% of total single nuclei (**Fig. 3J**), based on single-nuclei transcriptomic profiles (**Fig. 3K**).

These 15 cell clusters were then annotated to neural cell types based on the expression of neural marker genes in each cluster. A total of eight cell types were annotated (**Fig. 4A**). The majority of the neurons were annotated as GABAergic (GN) or glutamatergic (GluN) (**Fig. 4B**), the major neuron types present in human brain cerebral cortex. We next compared the snRNA-seq results across 9 experimental conditions (3 iPSC lines × 3 time points). To ensure that the same number of single nuclei were compared for each condition, random down-sampling was performed to match the single nuclei numbers in all 9 conditions (see **Fig. 4C**). We observed a significant increase in the number (or proportion) of glutamatergic neurons (GluNs) and a significant decrease in the number of GABAergic neurons (GNs) in the *FNDC4* KO iPSC-derived organoids when compared to WT organoids (see **Fig. 4D-4F**). This observation was consistent in two single-colony *FNDC4* KO iPSC-derived organoids at multiple time points. We repeated the random down-sampling three times, and each time observed the same result, suggesting that these results had not occurred by chance (**Fig. 4F** and **Supplemental Table S2**).

**Figure 4.**
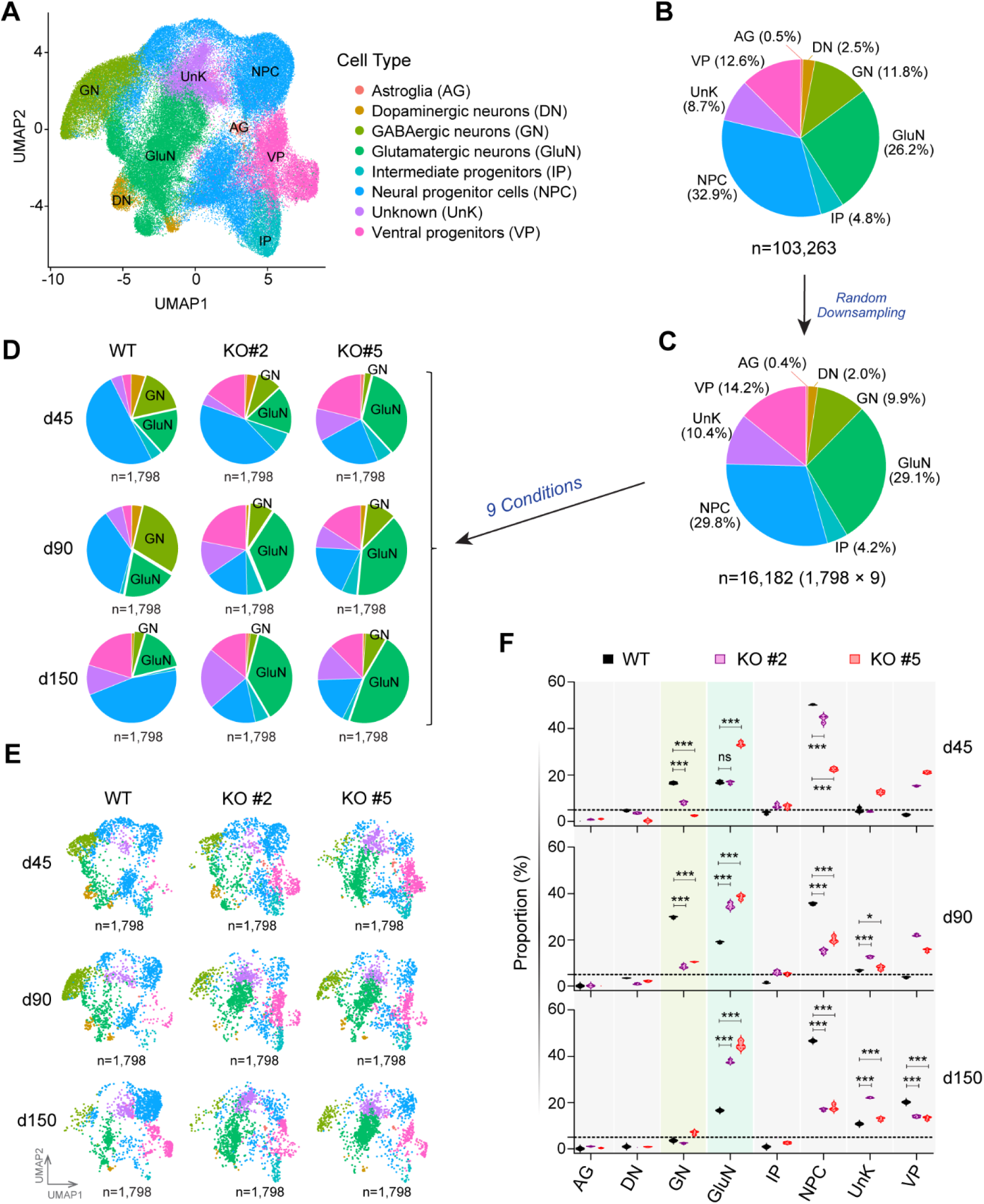
*FNDC4* KO led to an increased population of glutamatergic neurons defined by snRNA-seq. (**A**) Clustered single cells (n=103,263) were annotated to neural cell types based on the differential expression of marker genes. A total of 8 cell types were annotated and color-coded. (**B**) (**C**) Pie charts showing proportions of single cells for each annotated cell type in (**B**) total sequenced single nuclei (n=103,263), and in (**C**) random down-sampling single nuclei samples that included 9 experimental conditions (n=16,182). Random down-sampling was performed to ensure that each of the 9 experimental conditions included the same number of single nuclei (n=1,798) for comparison of cell type proportions. (**D**) Pie charts showing the proportions of annotated cell types in dorsal forebrain organoids for 9 individual experimental conditions (WT, KO#2 and KO#5 at day 45, 90, and 150 of organoid differentiation/maturation). (**E**) These single nuclei from 9 individual experimental conditions were also visualized by UMAP plots. (**F**) Violin plots comparing proportions of eight cell types in WT and two *FNDC4* KO organoids across three time points after three-rounds of random down-sampling. *P* values were calculated using two-way ANOVA with Dunnett’s multiple comparisons to WT samples. \**P* <0.05, \*\*\**P* <0.001, ns = not significant. *P* values for cell types with proportion of 5% (dashed line) or less were not shown.

Although a similar total number of single nuclei for each condition was used for single-nuclei RNA barcoding, the final numbers of single nuclei used for data analysis were variable across the 9 conditions studied after sequencing. To avoid the limitation of available single nuclei number (n=1,798) for the “WT-d150” condition, we removed the day-150 time point, thus making it possible to compare 7,305 single nuclei *per* condition (determined by “KO#5-d90”) across six conditions (**Supplemental Fig. S8C-F,** and **Supplemental Table S3**). We once again observed a significant increase in the number of GluNs and, simultaneously a decrease in the number of GNs in the *FNDC4* KO iPSC-derived organoids when compared to the WT organoids (**Supplemental Fig. S8D-F**). Taken together, these results demonstrated that *FNDC4* KO altered glutamatergic and GABAergic neuron proportions in these cerebral organoids in opposite directions.

### Transcriptomic profile of FNDC4-dependent neurogenesis

We next moved to explore potential molecular mechanism(s) involved in FNDC4-dependent neurogenesis. Based on all available single-nuclei transcriptomes (n=103,263), pseudo-time trajectories for neural development from less mature neural progenitor cells (NPCs) to more mature neurons could be generated (**Fig. 5A**). The pseudo-time trajectory analysis identified a “branch” point that led to distinct paths which generated GNs or GluNs (green cycle in **Fig. 5A**). When comparing the trajectories of WT and *FNDC4* KO organoids, this “branch” point could only be clearly identified in *FNDC4* KO but not in WT organoids (circled in **Fig. 5B**). Furthermore, this “branch” point overlapped a cell population of unknown cell type (UnK cells in **Fig. 5C**) for which the transcriptome profiles did not match any known neural cell types.

**Figure 5.**
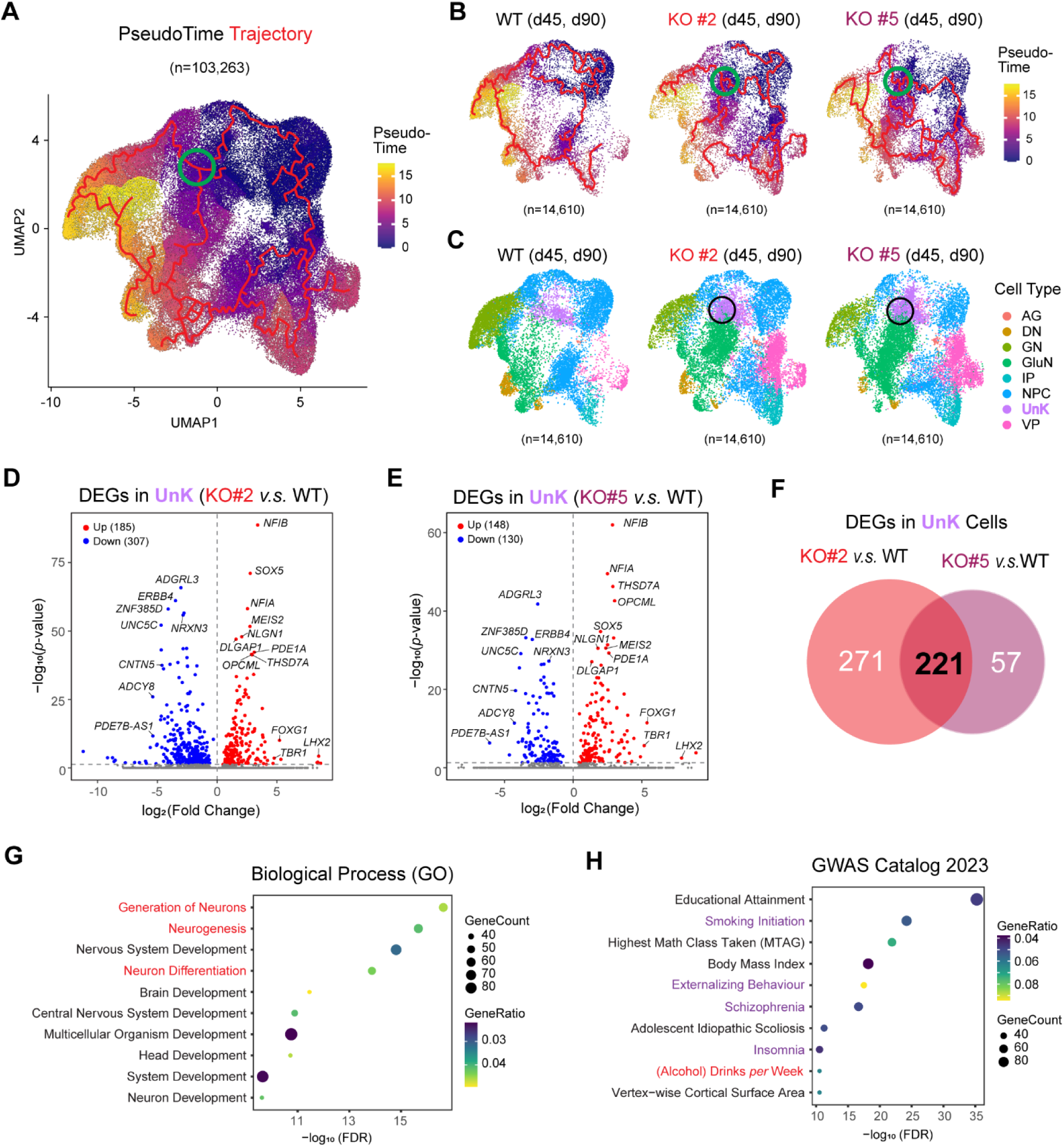
Transcriptome profile of FNDC4-dependent neurogenesis. (**A**) UMAP visualization of single-cell trajectory analysis inferring the order of these cells along neural developmental trajectories. Transcriptomic data for all single nuclei (n=103,263) from 9 experimental conditions were included in the analysis. Each cell/(dot) was assigned a pseudo-time value (color-coded) that represents which stage the cell is in along neural developmental trajectories. Cells at later stages of neural development (higher pseudo-time value) were colored in yellow. Trajectories for neural development were depicted as red lines. “Branch spots” leading to different neuronal subtypes were circled in green. (**B**) Single-cell trajectory analysis using single-nuclei transcriptomics of forebrain organoids differentiated from the WT (left), FNDC4 KO#2 (middle) and KO#5 (right) iPSC lines. Random down-sampling was performed to ensure that the same number of cells (n=14,610) from WT and KO organoids were included for comparison. Single nuclei of organoids differentiated from the same iPSC lines at two timepoints (d45, d90; n=7,305/timepoint) were combined to increase diversity (in developmental stages) and number of single cells for trajectory analysis. The “branch point” in the trajectories leading to different neuronal subtypes (circled in green) was observed in *FNDC4* KO but not in WT organoids. (**C**) UMAP visualization of the same single nuclei as in (b) with cell types color-coded. The “branch point” observed in *FNDC4* KO was overlapped with a cluster of “Unknown (UnK)” cells. (**D**) (**E**) Volcano plots for differentially expressed genes (DEGs) in “UnK” cells when comparing *FNDC4* KO to WT organoids. (**F**) Venn diagram showing 221 common DEGs in “UnK” cells were identified in organoids differentiated from both *FNDC4* KO iPSC lines. Top terms enriched by 221 DEGs in UnK cells after *FNDC4* KO in (**G**) the Gene Ontology (GO) Biological Process, and in (**H**) the GWAS Catalog phenotype/trait enrichment analysis. The false discovery rate (FDR) for pathway enrichment was computed from Fisher’s exact test and adjusted by using the Benjamini-Hochberg metho4d for the correction for multiple hypotheses testing.

These “UnK” cells might represent a cell type that is in “transition” in which cells have not yet committed to become either GN or GluN cells during the process of neural organoid differentiation. We next compared the transcriptomes of UnK cells from *FNDC4* KO to those in WT organoids in an effort to identify genes involved in the cellular “decision making” process for glutamatergic neurogenesis in the organoids after *FNDC4* KO. Differentially expressed genes (DEGs) with fold changes (FC) of more than 2 (|log_2_FC|≥1, adjusted *p* <0.05) in the UnK cells for each *FNDC4* KO organoid compared to WT were identified (**Fig. 5D, 5E**). A total of 221 DEGs, of which 107 were upregulated and 114 were downregulated, were consistently identified in both single-colony *FNDC4* KO organoids (**Fig. 5F** and **Supplemental Table S4**).

These *FNDC4*-dependent DEGs were used to perform Gene Ontology (GO)^34^ enrichment to help explore the biological function of FNDC4. The “top” enriched terms by GO Biological Process analysis were all related to neurogenesis (**Fig. 5G**), a result consistent with our observation of dysregulated GABAergic and glutamatergic neurogenesis in forebrain organoids after *FNDC4* KO (**Fig. 4**). The “top” DEGs (**Fig. 5D**, **5E**) included *NIFB*^35^, *FOXG1*^36, 37^, and *TBR1*^38^, all three of which have been reported to regulate GABAergic and/or glutamatergic neurogenesis. *FNDC4* KO resulted in a significant change in expression of these three genes during neural organoid differentiation. The GO Cellular Component and Molecular Function analyses have enriched terms such as “Cell Junction”, “Synaptic Membrane”, “Neuron to Neuron Synapse” and “Cell Adhesion Molecule Binding” (see **Supplemental Fig. S9**), suggesting a molecular function for FNDC4 that is involved in neural cell-cell interaction. Many of the “top” DEGs, including *ADGRL3*^39, 40^, *CNTN5*^41, 42^, *NRXN3*^43, 44^, *NLGN1*^45, 46^, and *OPCML*^47, 48^, encoded cell surface proteins with known roles in neurogenesis (**Fig. 5D, 5E**). These observations were consistent with the fact that FNDC4 is a transmembrane protein in the human brain (**Fig. 2G**) and that it has a functional domain (FN3) that may bind to protein partners and mediate cell-cell interactions^26^ (**Fig. 1D**).

Finally, the 221 *FNDC4*-dependent DEGs which we identified were also used for GWAS Catalog^49^ enrichment to help enhance our understanding of the potential role of the brain-expression of FNDC4 in clinical phenotypes/traits. Alcohol consumption (drinks *per* week) and multiple behavioral/neuropsychiatric phenotypes which often co-exist with AUD (smoking initiation, externalizing behavior, schizophrenia, and insomnia) were significantly enriched (**Fig. 5H**), supporting a possible functional role for brain-expression of FNDC4 in AUD. For example, genetic variants in “top” DEGs including *NRXN3*^50, 51^, *NFIA*^50^, *SOX5*^50^, *OPCML*^50, 51^, *ZNF385D*^52^, and *DLGAP1*^52^ (**Fig. 5D, 5E**), all of which have been associated with alcohol consumption^50, 51^ or sensitivity to alcohol^52^ by GWAS.

In summary, by taking advantages of snRNA-seq data generated using *FNDC4* KO iPSC-derived neural organoids, we were able to apply pseudo-time trajectory analysis of neural development and to identify a cell fate decision point leading to increased glutamatergic and decreased GABAergic neurogenesis in *FNDC4* KO organoids. Comparing the transcriptome profiles of those cells between WT and *FNDC4* KO, a series of DEGs were identified which suggested that *FNDC4* might be involved in AUD-related phenotypes, perhaps through a role in cell-cell interactions—interactions which are known to regulate neurogenesis^53–55^.

## DISCUSSION

In the present study, we characterized the function of a genome-wide significant SNP signal that was identified during a GWAS which we performed for AUD drug treatment response. That SNP proved to be an sQTL for the *FNDC4* gene in multiple human brain regions (**Fig. 1**). Our functional studies began with characterization of a GWAS SNP-associated *FNDC4* RNA splice isoform. We found that the FNDC4 splice isoform encoded a truncated protein that had lost the ability to localize to the membrane and be glycosylated, and which was rapidly degraded in the cytoplasm (**Fig. 2**), a result suggesting that the GWAS SNP might be associated with FNDC4 decrease or loss-of-function in the brain since FNDC4 protein in brain is a single-pass transmembrane glycoprotein (**Fig. 2**). We also characterized *FNDC4* function using CRISPR/cas9 gene editing, the creation of human iPSC-derived neural organoids and snRNA-seq (**Fig. 3**). We found that *FNDC4* knock-out resulted in a striking and significant shift in the relative proportions of glutamatergic and GABAergic neurons in iPSC-derived neural organoids (**Fig. 4**), suggesting a crucial role for FNDC4 in neurogenesis, perhaps through its function in mediating neural cell-cell interaction (**Fig. 5**). Our study characterized a novel genetic locus associated with AUD-related phenotypes, suggesting an FNDC4-dependent neurogenesis mechanism that contributes to AUD pathophysiology.

Our observation that *FNDC4* KO resulted in dysregulated GABAergic and glutamatergic neurogenesis (**Fig. 4**) is consistent with current understanding of AUD pathophysiology and the mechanism-of-action of drugs used to treat AUD. Alcohol directly interacts with GABA and glutamate receptors, and it is believed that AUD develops as a result of disrupted homeostasis of neurotransmission after exposure to alcohol^3^. AUD pharmacotherapy is also targeted to neurotransmission mechanisms. AUD often coexists with other substance use disorders, particulaly smoking, and with mental health disorders^2^. Our data support the possibility that *FNDC4* might be a pleiotropic gene which contributes to additional neuropsychiatric disorders. Specifically, those FNDC4-dependent DEGs have been reported to enrich “smoking initiation” and “schizophrenia” as clinical phenotypes (**Fig. 5H**), and several “top” FNDC4-dependent DEGs are known risk genes for different psychiatric disorders. For example, the *THSD7A* gene, for which expression is significantly increased after *FNDC4* KO (**Fig. 5D, E**), has been identified in GWAS for bipolar disorder risk^56, 57^ and anticonvulsants treatment response^58^. In addition, a genetic locus (index SNP, rs12474906) which is an expression QTL for *FNDC4*^14^ has been identified by large-cohort GWAS as a pleiotropic locus for eight different psychiatric disorders (*p* = 8×10^-9^) including schizophrenia, bipolar disorder, and attention-deficit/hyperactivity disorder^59^, all of which are highly comorbid with AUD^60–64^. The fact that *FNDC4* KO leads to increased glutamatergic but decreased GABAergic neurogenesis is also compatible with observations made in *Fndc4* KO mice which display hyperactivity (increased vertical activity and limb grasping)^65^. Hyperactive behavior was found to be related to an imbalance between inhibitory (GABAergic) and excitatory (glutamatergic) neuron development^66^ as well as increased glutamatergic transmission^67, 68^. In addition, childhood attention-deficit/hyperactivity disorder and externalizing behavior are significant risk factors for the development of alcohol and substance use disorders^69–71^.

Our results suggest a possible scenario in which individual adults might have differing ratios of GABAergic and glutamatergic neurons, and those interindividual differences in neuron populations might be, at least in part, due to genetic polymorphisms related to the *FNDC4* gene which are linked to individual variation in AUD risk and drug treatment response. Specifically, our GWAS showed that variant SNP alleles that were associated with increased FNDC4 splicing isoforms (**Fig. 1B**) which encode a loss-of-function FNDC4 truncated protein (**Fig. 2**) were also associated with adverse drug treatment response (**Fig. 1A**). Since FNDC4 loss-of-function results in increased glutamatergic but decreased GABAergic neurogenesis (**Fig. 4**), our results also raise the possibility that AUD patients who carry the variant SNP allele (with increased expression of the FNDC4 splicing isoform) might benefit from therapy that antagonizes glutamatergic neurotransmission. Our results may also provide a genetic basis for individualized AUD treatment.

Finally, we should point out the limitations of our studies, beginning with our use of *in vitro* CNS models. We used human iPSC-derived dorsal forebrain organoids to study FNDC4 molecular function because FNDC4 is expressed more highly in human brain cerebral cortex than in other brain regions (GTEx)^14^. To our knowledge, FNDC4 function in human CNS model(s) has not been reported previously. However, FNDC4 is also expressed in other brain regions. Whether it functions in the regulation of neurogenesis in other brain regions clearly needs to be studied further. Although iPSC-derived neural organoids provide an opportunity to manipulate human gene expression in brain-like models and to capture neural cell-cell interactions during neural development, this model obviously does not fully capture neural function in the adult human brain. Therefore, FNDC4 neural functions other than neurogenesis would not necessarily be captured in our study using neural organoids. In addition to genetics, environmental factors include alcohol exposure and their interaction with genetics might also play a critical role in risk for developing AUD. Whether environmental factors might affect FNDC4 function in the brain should be studied further. Animal studies will also be required to investigate the role of *Fndc4* in addiction behavior. Although behavioral changes in hyperactivity, a risk factor for developing AUD, have already been observed in *Fndc4* KO mice, whether those *Fndc4* KO mice will develop alcohol-related behaviors is currently unknown and needs to be studied. Those animal models might also provide an opportunity to test novel therapeutic interventions targeting FNDC4-related mechanism in future studies.

In summary, we have studied the function of the *FNDC4* gene for which sQTLs in the brain have been associated with AUD risk and variation in drug treatment response^13^. We demonstrated that the *FNDC4* sQTL related mRNA splice isoform results in loss-of-function of *FNDC4* which then can lead to the dysregulation of the “balance” between glutamatergic and GABAergic neurogenesis. These results strongly support the possibility that FNDC4 function in the CNS might represent one factor playing a role in AUD risk and variation in response to the drug treatment of AUD, providing a genetic basis for individualized AUD pharmacotherapy and an opportunity for the development of potential novel AUD therapies.

## METHODS

### Cell culture

HEK293T cell lines were obtained from the American Type Culture Collection (ATCC). HEK293T cell lines were cultured in Dulbecco’s Modified Eagle Medium (DMEM) with 10% FBS. A human episomal iPSC line was obtained from ThermoFisher (Cat#: A18945). After directed differentiation and teratoma analyses, the iPSCs retained their differentiation potential for the ectodermal, endodermal, and mesodermal lineages. All cells tested negative for virus including HIV, HTLV, HSV, CMV, EBV, HBV and HCV. Cytogenetic analysis demonstrated an apparently normal female karyotype. Human iPSCs were cultured in StemFlex™ medium (Cat#: A3349401) following the manufacture’s protocol.

### FNDC4 overexpression

The human *FNDC4* cDNA plasmid construct was purchased from OriGene (Cat#: RC207471). This construct overexpresses a canonical FNDC4 protein fused with a MYC-FLAG-tag at the C-terminus. The *FNDC4* splice variant cDNA was synthesized by GenScript and was cloned into the same vector (pCMV6-Entry) as the *FNDC4* cDNA plasmid purchased from OriGene. HEK-293T cells were transfected with *FNDC4* cDNA constructs by using Lipofectamine™ 3000 Transfection Reagent (ThermoFisher). Eight hours after transfection, cell culture media was replaced with fresh media. After ∼48 hours of transfection, cells were harvested for protein extraction using Pierce™ IP Lysis buffer (Cat#:87787). Protein concentrations were determined by using the Pierce™ BCA Protein Assay Kit (Cat#:23227).

### Animals

Male Wistar rats (350-375g) were purchased from Charles River Laboratories (Wilmington, MA). The tissues from the cortical and subcortical brain regions included caudate putamen (CPU), nucleus accumbens (NAC), amygdala (AMG), lateral orbitofrontal cortex (lOFC), infralimbic/prelimbic cortices (IL/PL), anterior cingulate cortex (ACC), and “extra’” forebrain regions including medial orbitofrontal and motor cortices (Extra) that were harvested from three rats. Immediately after harvesting, all tissues were kept on dry ice and then stored in a -80°C freezer until further processing. For protein lysates preparation, those rat brain tissues were exposed to IP lysis buffer at a w/v ratio of 1mg/10µL followed by homogenization using a ½ mL syringe (BD; Cat#:305620). Protein concentrations were determined by BCA assay. Animal tissues were harvested in accordance with the U.S. National Institutes of Health Guidelines for the Care and Use of Laboratory Animals and approved by the Mayo Clinic Institutional Animal Care and Use Committees (IACUC).

### Western Blot Analysis

Denatured proteins were loaded onto a 4–20% Mini-PROTEAN® TGX™ Precast Protein Gels (Bio-Rad) to separate proteins. Precision Plus Protein Dual Color Standards (Bio-Rad; Cat#:1610374) were used as protein markers for all blots included in this study. Proteins were transferred from the gels to PVDF membranes which were then blocked with 5% non-fat milk at room temperature for 1 hour. After washing with TBST, membranes were incubated with primary antibody (see **Supplemental Table S1** for detailed information), which was dissolved in 1% BSA prepared in TBST at 4°C overnight with gentle rocking. Following incubation, the membranes were washed vigorously three times in TBST buffer and were then incubated with horseradish peroxidase (HRP)-labelled secondary antibody, which was dissolved in 5% non-fat milk at room temperature for 1 hour. The SuperSignal West Dura Extended Duration Substrate (ThermoFisher, Cat#:34075), a luminol-based enhanced chemiluminescence HRP substrate, was applied to the membranes, and radiographic images were captured by use of the ChemiDoc™ Touch Image System (Bio-Rad).

### Subcellular Fractionation

Membrane, cytoplasmic and nuclear protein fractions were prepared using the Subcellular Protein Fractionation Kit for Cultured Cells (ThermoFisher; cat#:78840) following the manufacture’s protocol. Briefly, HEK293T cells with FNDC4 overexpression were collected and rinsed with cold PBS. Cell pellets were immersed in the first reagent which causes selective cytoplasmic membrane permeabilization, releasing soluble cytoplasmic contents. After collecting the cytoplasmic contents, a second reagent was added to dissolves cytoplasmic, endoplasmic, and mitochondria membranes but which did not solubilize nuclear membranes. After centrifugation, the intact nuclei were pelleted, and the supernatants were collected as was the membrane fraction. Nuclear pellets were used for soluble nuclear protein extraction. This protocol can also further separate soluble and chromatin-bound nuclear proteins, a step which was not performed in our study.

### Protein Deglycosylation

The PNGase F (Cat#: P0704S) protocol from New England Biolabs was used for the protein deglycosylation assay. Briefly, 10 µg of protein lysate from HEK293T cells with FNDC4 overexpression were denatured by heating at 100°C for 10 minutes. Denatured proteins were mixed with 1× GlycoBuffer 2 and 1% NP-40 to a total reaction volume of 20 µL. After adding 1 µL PNGase F, the reaction was incubated at 37°C for 1 hour. Results of deglycosylation were assessed by Western blot assay.

### FNDC4 KO by CRISPR/Cas9

The Alt-R™ CRISPR/Cas9 System (IDT) was used for *FNDC4* gene editing as described in our previous studies^72^. This system includes a Hi-Fi Cas9 endonuclease, a gene-specific crRNA and a universal tracrRNA. These three components were combined to form a CRISPR/Cas9 ribonucleoprotein (RNP) complex and the RNP complex was delivered into cells for target DNA sequence cutting. Two crRNAs targeting *FNDC4* exon 3 and 4 DNA sequences (see **Supplemental Table S1** for crRNA sequences), respectively, were designed for “double cuts” which would allow single-colony selection using standard PCR and agarose gel electrophoresis. The CRISPR/Cas9 RNP complex was delivered into iPSCs by electroporation using the P3 Primary Cell 4D-Nucleofector™ X Kit (Cat#: V4XP-3032, Lonza) according to the manufacturer’s instructions (Pulse code: CM-113). Two days after electroporation, edited iPSCs were passaged to form single colonies. A portion of the cells (10%) was lysed with DNAzol® Direct (DN 131, Molecular Research Center) and the lysates were used as PCR templates for the evaluation of CRISPR/Cas9 editing efficiency. The sequences of primers for amplification of the edited sites are listed in the Major Recourse table. The PCR was conducted with the KAPA HiFi HotStart ReadyMix PCR Kit (Cat#: KK2601, Roche). Once single colonies were formed, they were genotyped by PCR for the selection of homozygous KO colonies. Selected single-colony KO iPSCs were expanded for two more passages and were genotyped once again to confirm their KO status. Confirmed single-colony KO iPSCs were further characterized by karyotyping to ensure genomic integrity and by trilineage differentiation to ensure their pluripotency.

### Generation of Forebrain Organoids from iPSCs

The STEMdiff™ Dorsal Forebrain Organoid Differentiation Kit (Cat#: 08620, STEMCELL Technologies) was used to generate forebrain region-specific organoids by following the manufacture’s protocol. That protocol can reliably and reproducibly generate forebrain organoids with cell types appropriate for the human cerebral cortex^31, 73^. Briefly, WT and *FNDC4* KO iPSCs were seeded in an AggreWell™800 24-well plate for uniform embryoid bodies (EBs) formation. After 6 days, the EBs were transferred to a 6-well ultra-low adherent plate (∼30 EB aggregates *per* well) and were placed on a level shaker in the 37°C incubator for expansion until day 25. Expanded organoids were further cultured for ∼20 days of forebrain organoid differentiation. Differentiated forebrain organoids were cultured in maintenance media until used. Specifically, three organoids differentiated from each iPSC lines were harvested at days 45, 90 and 150 for single nuclei isolation and fixation which were also used for snRNA-seq. At day 150, organoids were also harvested for cryosection and immunostaining.

### Cryosection and Immunofluorescence (IF)

Neural organoid cryosection and immunofluorescence were performed following the protocol entitled “Cryogenic Tissue Processing and Section Immunofluorescence of Neural Organoids” from STEMCELL. Specifically, forebrain organoids were fixed with 4% paraformaldehyde (PFA) overnight at 4°C. After fixation, organoids were equilibrated in 30% sucrose overnight at 4°C and were then snap frozen and transferred to a -80°C freezer until used. Sectioning was performed in a Leica CM1520 cryostat at -26°C with the section thickness set at 16 μm. Multiple serial sections were collected and mounted on glass slides. Sectioned slides were kept in a -20°C freezer until used. Before immunostaining, sectioned slides were removed from the freezer and were allowed to dry at room temperature. Sections were then outlined with a Pap pen and were washed with PBST for 10 mins at 37°C to remove gelatin. Sections were blocked with 5% BSA/PBST for 1 hour, followed by primary antibody incubation overnight at room temperature in a humidified chamber. After 3 washes with PBST, sections were incubated with fluorescence-labelled secondary antibodies at room temperature for 2 hours. After an additional 3 washes with PBST, slides were air dried at room temperature. Slides were then mounted with Mountant and were covered with coverslips. Stained slides were stored at 4°C before imaging using a Zeiss LSM 780 Confocal Microscope. See **Supplemental Table S1** for antibody information including vendor’s catalog numbers and dilution factors.

### Single Nuclei Isolation and Fixation

Single nuclei were isolated from fresh neural organoids harvested at days 45, 90 and 150 of differentiation/maturation. Specifically, three organoids generated from each iPSC line were transferred into a 15-mL tube and were rinsed with 1 mL of cold DPBS. After removing DPBS, chilled Lysis Buffer (Tris-HCl, 10 mM; NaCl 10nM; MgCl_2_, 3mM; NP-40, 0.1%) was added and samples were incubated on ice for 30 mins to lyse the organoids. Organoids were then further triturated by pipetting 5-7 times to release nuclei. Nuclei were then pelleted by centrifugation at 500×g for 10min at 4°C. Nuclear pellets were resuspended in Nuclei Wash and Resuspension buffer (1% BSA and 0.2U/µL RNase inhibitor in PBS) and were mixed by gentle pipetting. Lysis efficiency was assessed by trypan blue staining with automated cell counting. If a high cell viability (>10%) remained, the lysis process was repeated until cell viability fell below 10%. Cell debris and large clumps were removed by using a 40 μm Flowmi Cell Strainer. Filtered single nuclei were washed twice using the Nuclei Wash and Resuspension buffer and were centrifuged at 500×g for 10 min at 4°C. Washed single nuclei were directly fixed using the Evercode™ Nuclei Fixation v2 kit (Parse Biosciences) by following the manufacture’s protocol. Fixed and permeabilized nuclei were snap frozen and stored at -80°C until used.

### snRNA-seq

Single-nucleus RNA samples were barcoded using the “split-pool” combinatorial barcoding technology^33^. Specifically, frozen single nuclei were thawed on ice. Single nuclei numbers and viability were analyzed using Nexcelom Cellometer Auto2000 with the AOPI fluorescent staining method. Single-nuclei library preparation was performed using the Evercode™ WT v2 kit (Parse Bioscience) according to the manufacturer’s protocol. Approximately 16,000 fixed and permeabilized nuclei *per* sample were loaded into 48 wells of the Round 1 Plate. RNA was reverse transcribed using oligo dT and random hexamer primers with a well-specific barcode that was associated with specific samples. After 3 rounds of combinatorial barcoding, a total of ∼150,000 nuclei from 9 samples were recovered. Barcoded nuclei were then split into 8-sublibrary tubes (Round 4) and were lysed (see **Fig. 3A**). After nuclear lysis, cDNA was captured, amplified, and quantified by Qubit DNA HS assay kit. The multiplexed sublibraries were pooled and sequenced on a Novaseq X Plus, 10B flowcell using paired-end 150nt (PE150) mode. Library preparation and sequencing were performed at the Northwestern University NUSeq core.

### snRNA-seq Data Analysis

The raw sequencing FASTQ data were processed using the Parse Bioscience split-pipe pipeline (v1.4.0) with default settings to align sequencing reads to the human genome (hg38) and to demultiplex samples. Downstream analysis was performed in R (v 4.3.2), with data filtering and initial analyses performed using Seurat (v.5.0.3)^74^. Genes were filtered to remove those expressing in less than 100 nuclei, and nuclei were filtered to only include those with at least 500 genes detected. A set of 2000 genes with the highest variance across all cells were then chosen as variable features. All 9 single-nuclei forebrain organoid samples (3 iPSC lines × 3 time points) were then integrated together, and anchor genes were identified for further unsupervised clustering analyses. Clustering of single nuclei was performed using the Uniform Manifold Approximation and Projection (UMAP)^75^, identifying 15 distinct single nuclei clusters with a resolution of 0.5. Clusters were then labeled for distinct brain cell types using ScType^76^ with prior neural organoid datasets^30, 77^. Cluster marker genes were determined using the Seurat function FindAllMarkers for each cluster and filtered by Bonferroni-corrected *p* value < 0.05 and |log2 fold change| > 1. We compared the difference between conditions by looking at the clustering differences, differential gene expression between samples, as well as differential gene expression between clusters of the same samples. When comparisons between samples were made, random down-sampling was performed using the “sample()” function on the larger sample to allow for equal number of nuclei/cells for major comparisons, and multiple down-samples were used.

Because these forebrain organoids included cells at various differentiation and maturation states, trajectory and pseudotime analyses were performed using the Monocle3 package (v. 1.3.7)^78^, with trajectories calculated independently between samples. For all samples pooled, a resolution of 3e-5 was used, while for WT alone, KOc2 alone, and KOc5 alone (all including both d45 and d90), a resolution of 3e-3 was used. A root cluster was set in the Neural Progenitor Cell cluster(s) for all samples from which pseudotime and trajectories could branch.

## DATA AVAILABILITY

The snRNA-seq data generated in this study have been deposited in NCBI’s Gene Expression Omnibus (GEO) and are accessible through GEO Series accession number: GSE285126.

## AUTHOR CONTRIBUTIONS

XZ, DL and RMW designed the study. XZ, LW, SK, ED, AS, AJ, CMW, IM, TMK, and DL performed the experiments and collected the data. XZ, AJJ, IMG, HG, and DL analyzed the data. XZ, AJJ, RMW, and DL wrote the manuscript. BJC, NS, LieW, MAF, JMB, VK, and HL contributed to the interpretation of the results and revised the manuscript. DL and RMW supervised this project. All authors read and approved the final version of the manuscript.

## ACKNOWLEDGMENTS

This work was supported in part by National Institutes of Health (NIH) grants R01 GM028157, R01 AA027486, and by the Mayo Clinic Center for Individualized Medicine (CIM). S. Kim was supported by the NIH T32 Training Grant in Clinical Pharmacology (GM08685). We want to thank the GTEx Project for generating the datasets that have been cited in this study.

