## Supplemental Figures S1-S9 for "Alcohol Use Disorder Associated Gene *FNDC4* Alters Glutamatergic and GABAergic Neurogenesis"

**Figure S1.** The rs56951679 SNP (chr2:28260851; T>C) is an sQTL for FNDC4 in multiple human brain regions (Cortex, Nucleus Accumbens, Frontal Cortex, Hippocampus and Caudate) based on the RNA-seq data generated by GTEx. The variant allele, “C”, is associated with an increased level of intron excision at chromosome 2: 27,492,478-27,493,389 (human genome assembly build 38).

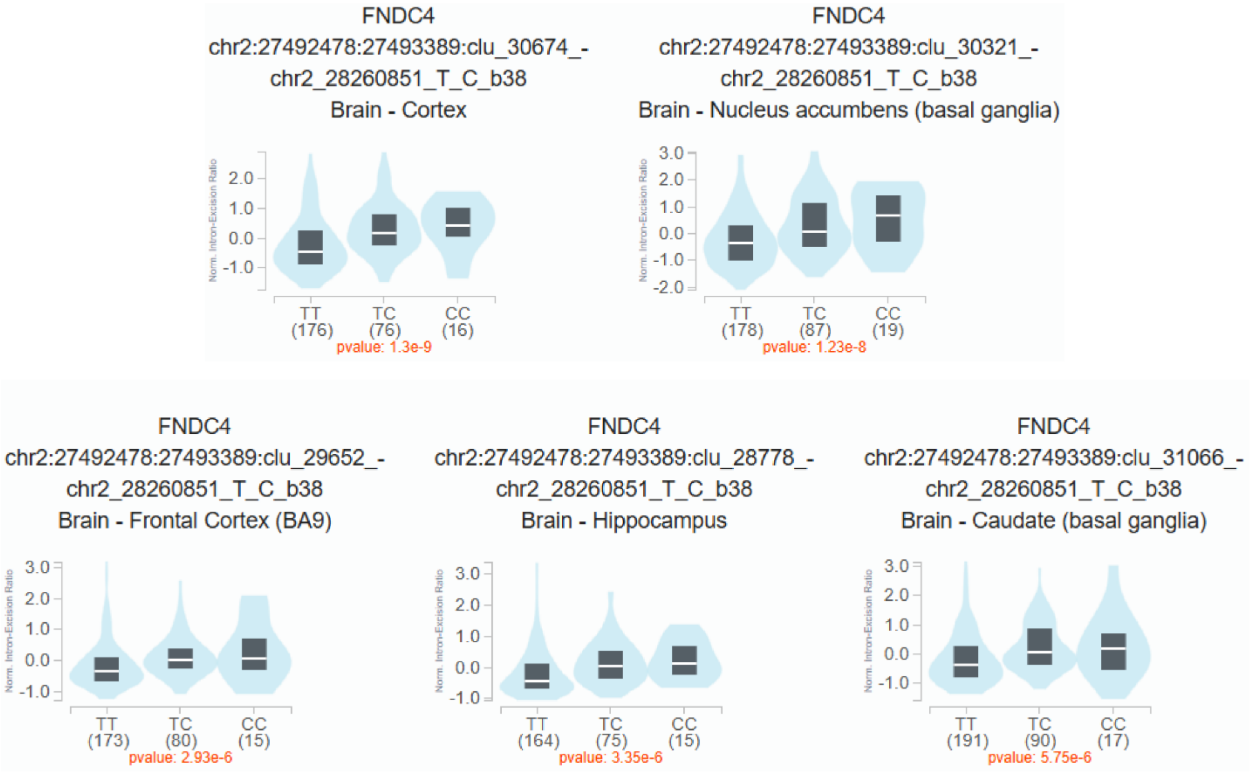

**Figure S2.** The rs1260326 SNP (chr2:27508073; C>T), which has been associated with AUD and alcohol consumption in GWAS, is an sQTL for FNDC4 in multiple human brain regions based on the RNA-seq data generated by GTEx. The common allele "C" is associated with an increased level of intron excision at chromosome 2: 27,492,478-27,493,389 (human genome assembly build 38).

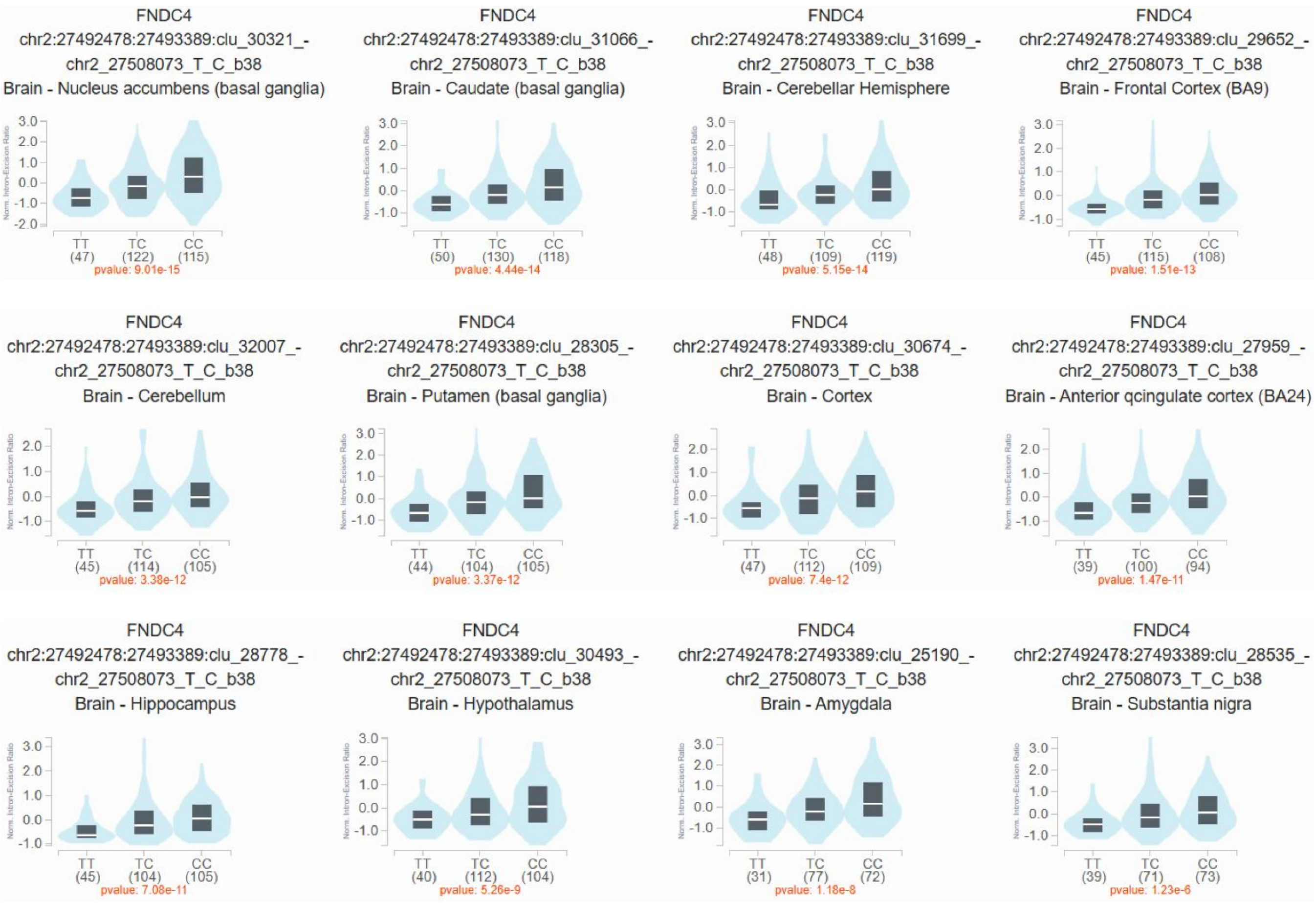

**Figure S3.** (**A**) The Western blot shows cDNA-overexpressed canonical and truncated FNDC4 proteins (fused with MYC- and FLAG-tags at their C-termini) using anti-MYC-tag antibodies. Beta-actin (ACTB) was blotted as a loading control. (**B**) Those same protein samples were then used to test anti-FNDC4 antibodies. Five of those tested FNDC4 antibodies detected overexpressed FNDC4 proteins, but they varied in efficiency and specificity. (**C**) Table shows the basic information of 5 tested FNDC4 antibodies (see Table S1 for details). Antigen aa number was based on the aa sequences of canonical FNDC4. (**D**) Western blots for endogenous FNDC4 presents in protein lysates of human whole brain (WB), cerebral cortex (CC) and cerebellum (CB). Overexpressed canonical and truncated FNDC4 proteins were blotted as positive controls. Protein marker (M) was loaded to separate human brain protein lysates from overexpressed FNDC4 proteins. (**E**) Detection of FNDC4 proteins in culture media for HEK293T cells overexpressing FNDC4 proteins (fused with MYC- and FLAG-tags at their C-termini) using anti-MYC-tag antibody. Culture media collected in three overexpression experiments (Exp) were concentrated (20 ×) and tested with cell lysate in a same blot. Protein marker (M) was loaded to separate cell lysate and culture media samples.

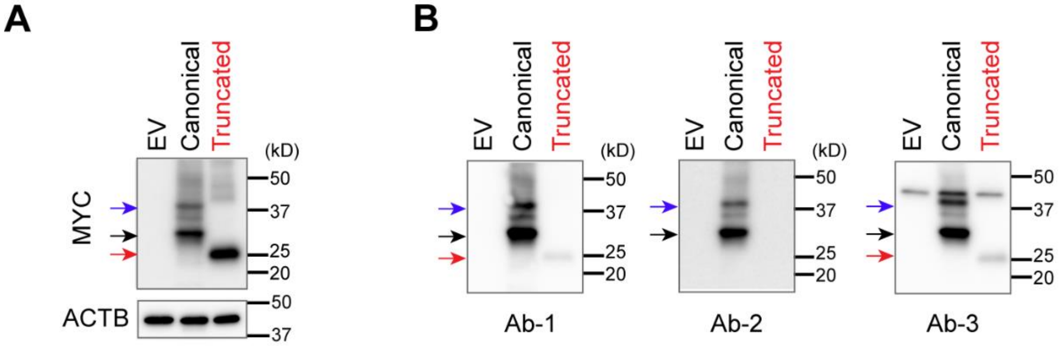


**C**

| FNDC4 Ab# | Clone ID | Antigen amino acids (aa) |
|---|---|---|
| Ab-1 | OTI3B11 | aa1 – aa234 |
| Ab-2 | N/A | aa194 – aa234 |
| Ab-3 | OTI1B7 | aa1 – aa234 |
| **Ab-4** | **OTI3E10** | aa1 – aa234 |
| Ab-5 | 7F9 | aa45 – aa154 |

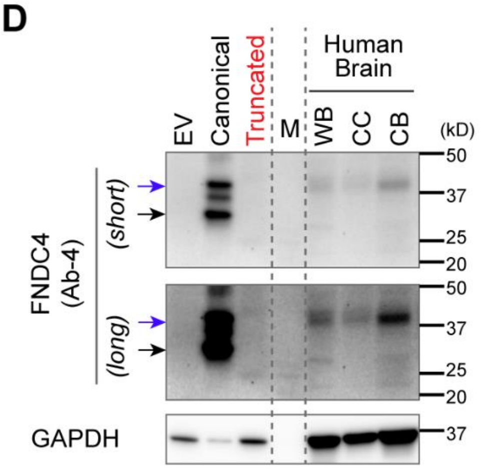


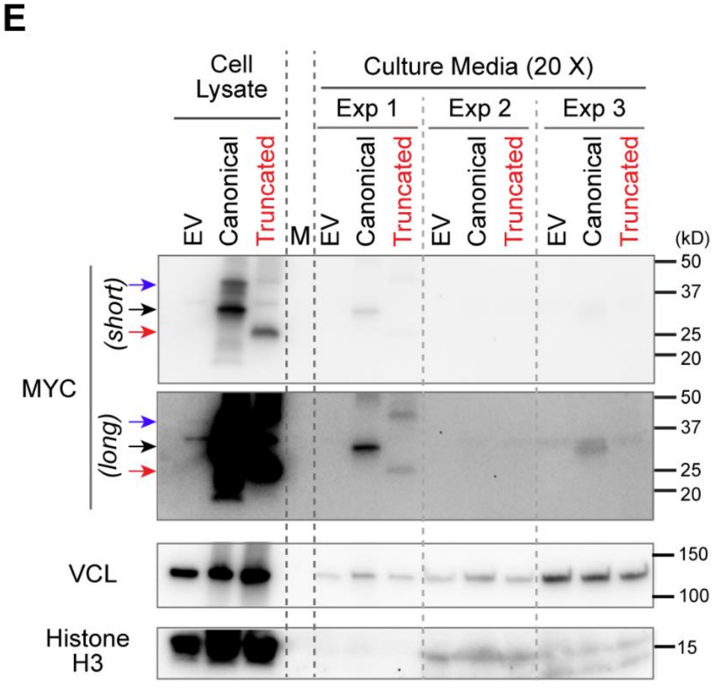

**Figure S4.** Amino acid sequences alignment for human, rat and mouse FNDC4 proteins. Signal peptide (SP) sequences were highlighted in red. Alignment was performed using the protein sequence data in the UniProtKB.

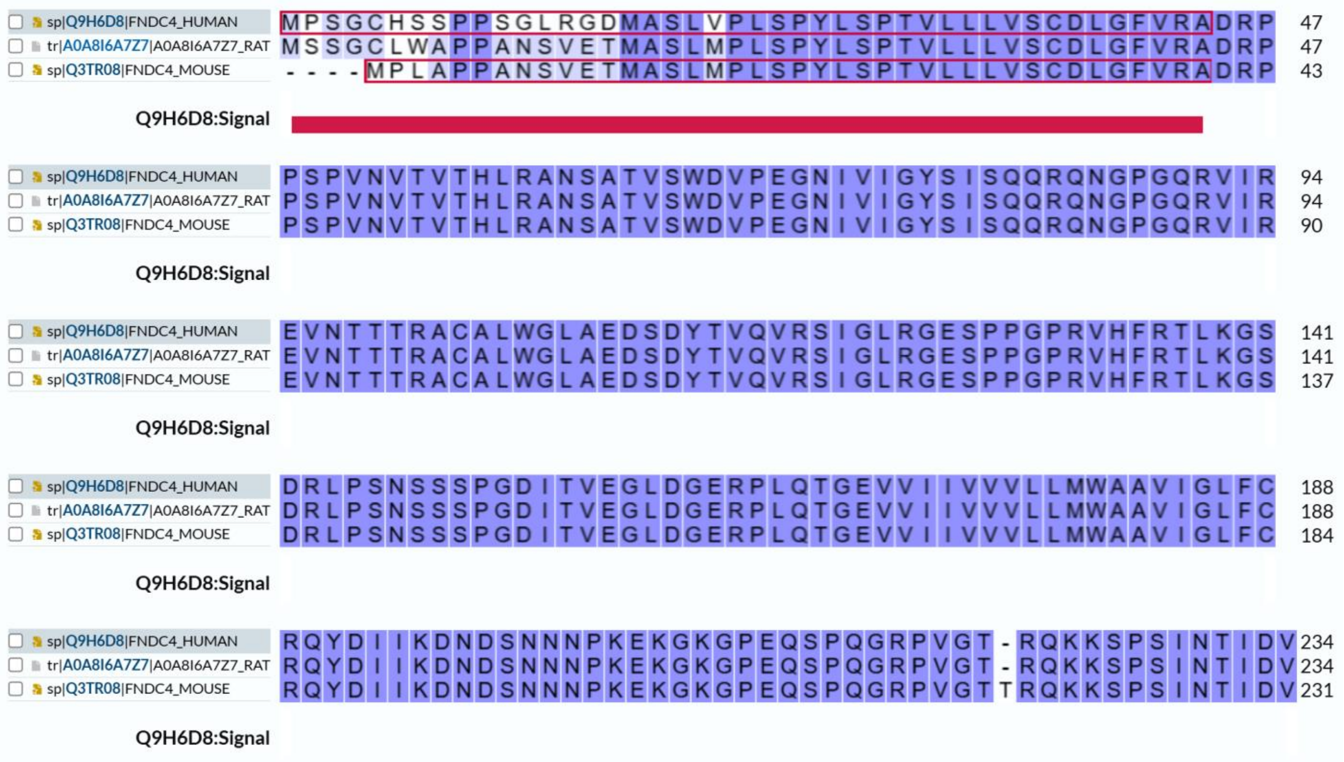

**Figure S5. Generation and Characterization of *FNDC4* KO iPSCs**. (**A**) Scheme for the generation of *FNDC4* KO iPSC lines using CRISPR/cas9 gene editing. (**B**) Design of guide RNAs (gRNAs) for *FNDC4* KO, and of PCR primers for KO colony selection. Successful CRISPR/cas9 editing by both gRNAs has been designed to remove 388 base pairs (bp) from the *FNDC4* gene, resulting in a 554 bp amplicon by PCR using designed primers. (**C**) Agarose gel results showing the PCR amplicons when genomic DNAs from the WT control and CRISPR/cas9-edited iPSCs (mixed) were used as PCR templates. The presence of a smaller band (~554 bp) in the edited iPSCs (lanes 1 and 2) indicates successful *FNDC4* KO in certain individual cells. Lanes 1 and 2 are the same CRISPR/cas9 experiments with slight differences in transfection methods to deliver CRISPR/cas9 RNP complexes. These "mixed" *FNDC4* KO iPSCs were used for single-colony isolation. (**D**) Selection of single-colony *FNDC4* KO iPSCs by PCR. Genomic DNA extracted from single-colony iPSC lines was used as PCR templates and the PCR amplicons were visualized in agarose gel. Four single-colony iPSC lines (colony numbers highlighted in red) which are potentially homozygous *FNDC4* KO were passaged and populated for further validation. (**E**) After 3 passages of the four *FNDC4* KO iPSC lines, they were genotyped once again by PCR to validate the homozygosity of *FNDC4* KO. Colonies A2 and A5 (renamed as colonies #2 and #5, respectively), which maintained the homozygosity of *FNDC4* KO, were used in further studies. (**F**) Both of the homozygous *FNDC4* KO iPSC lines had their genomic integrity confirmed by karyotyping.

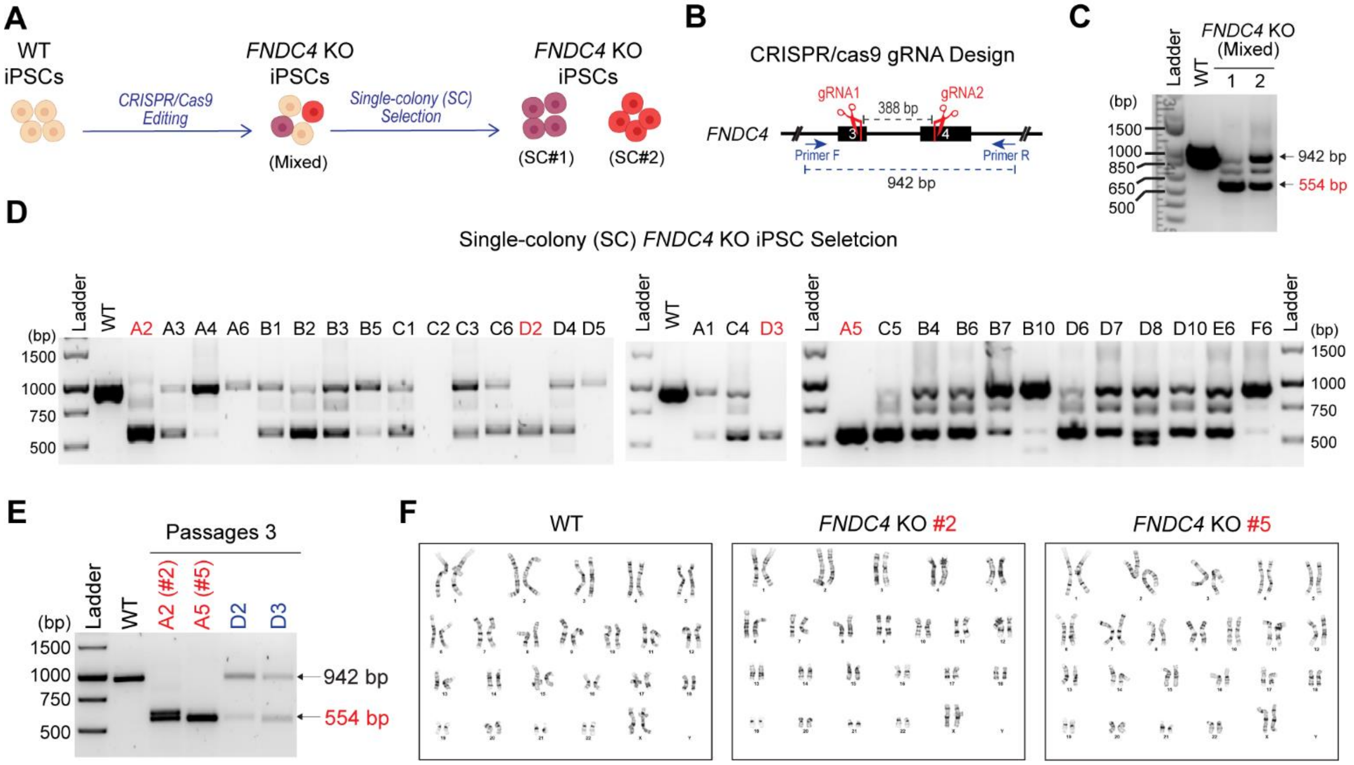

**Figure S6. Generation and Characterization of forebrain organoids from human iPSCs**. (**A**) Scheme for forebrain organoid generation. Human iPSCs were seeded in AggreWell™800 to form embryoid bodies (EBs). After 6 days, EBs were transferred to a suspension culture plate for organoid expansion until day 25. Organoids were then cultured in forebrain organoid differentiation medium. After approximately 3 weeks of differentiation, organoids were switched to maintenance media and cultured for up to 150 days. Three organoids from each iPSC line (WT, *FNDC4* KO#2 or KO#5) were harvested at three time points (d45, d90 and d150) for single nuclei isolation and snRNA-seq. Forebrain organoids at day 150 were cryosectioned and characterized by immunofluorescence (IF) staining of neural markers. IF of (**B**) neural progenitors (PAX6) and neuron (MAP2) markers, (**C**) superficial (SATB2) and deep (BCL11B, also known as CTIP2) neuron markers, (**D**) developing forebrain neuron markers (MAP2 and FOXG1), and (**E**) glutamatergic (SLC17A7, also known as VGLUT1) and GABAergic (GAD1, also known as GAD-67) neuron markers.

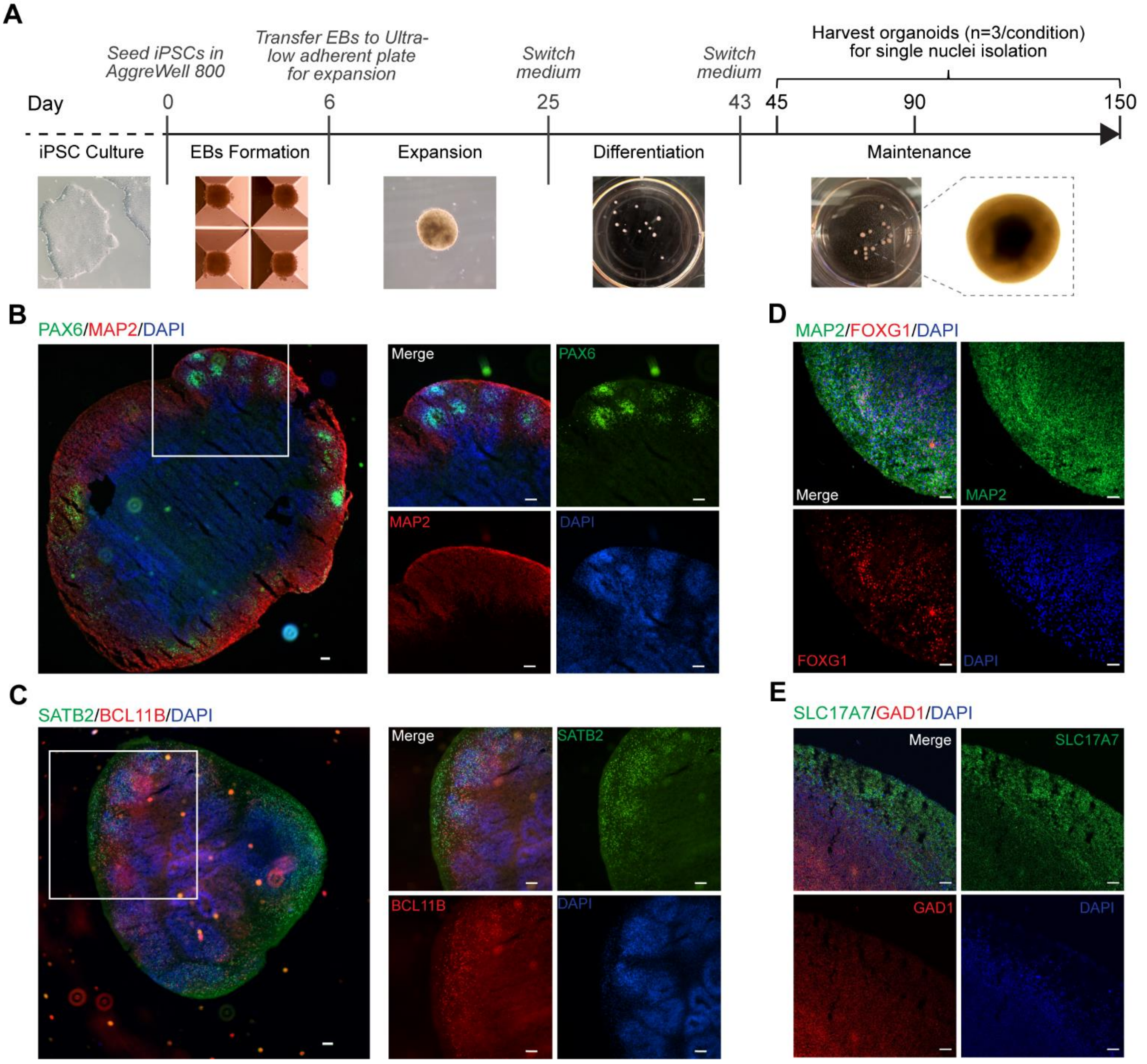

**Figure S7.** Bubble plot showing differentially expressed marker genes for cell type annotation. Each line is a gene marker, and each column is an annotated cell type. NPC, neural progenitor cells; GluN, glutamatergic neurons; VP, ventral progenitor; GN, GABAergic neurons; AG, astroglia; IP, intermediate progenitor; DN, dopaminergic neurons; UnK, unknown.

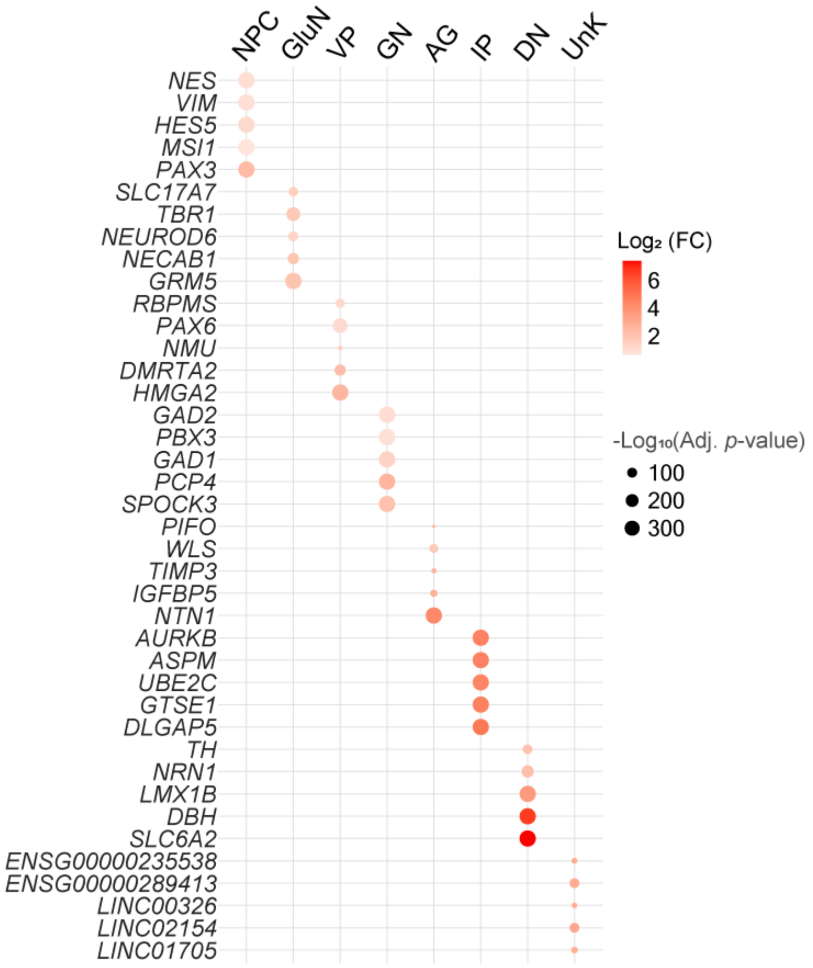

**Figure S8.** Pie charts showing the proportion of single nuclei for each annotated cell type in (**A**) total sequenced single nuclei (n=103,263), and in random down-sampling single nuclei samples under (**B**) 9 or (**C**) 6 experimental conditions. Random down-sampling was performed to ensure that each experimental condition would include a same number of single nuclei for comparison of cell type proportions. That same number was determined by single-nuclei number of the condition which had the least available single nuclei. (**D**) Pie charts showing the proportions of annotated cell types in dorsal forebrain organoids for 6 individual experimental conditions (WT, KO#2 and KO#5 at day 45, and 90 of organoid differentiation/maturation). When pie charts for individual conditions were compared, increases in the proportion of GluN (green) were observed in *FNDC4* KO organoids. (**E**) These single nuclei from 6 individual experimental conditions were also visualized by UMAP plots. (**F**) Violin plots comparing proportions of eight cell types in WT and two *FNDC4* KO organoids at days 45 (d45) and 90 (d90) after three-rounds of random down-sampling. *P* values were calculated by two-way ANOVA with Dunnett's multiple comparisons to WT samples. **P* <0.05, ****P* <0.001, ns = not significant. *P* values for cell types with a proportion of 5% (dashed line) or less were not shown.

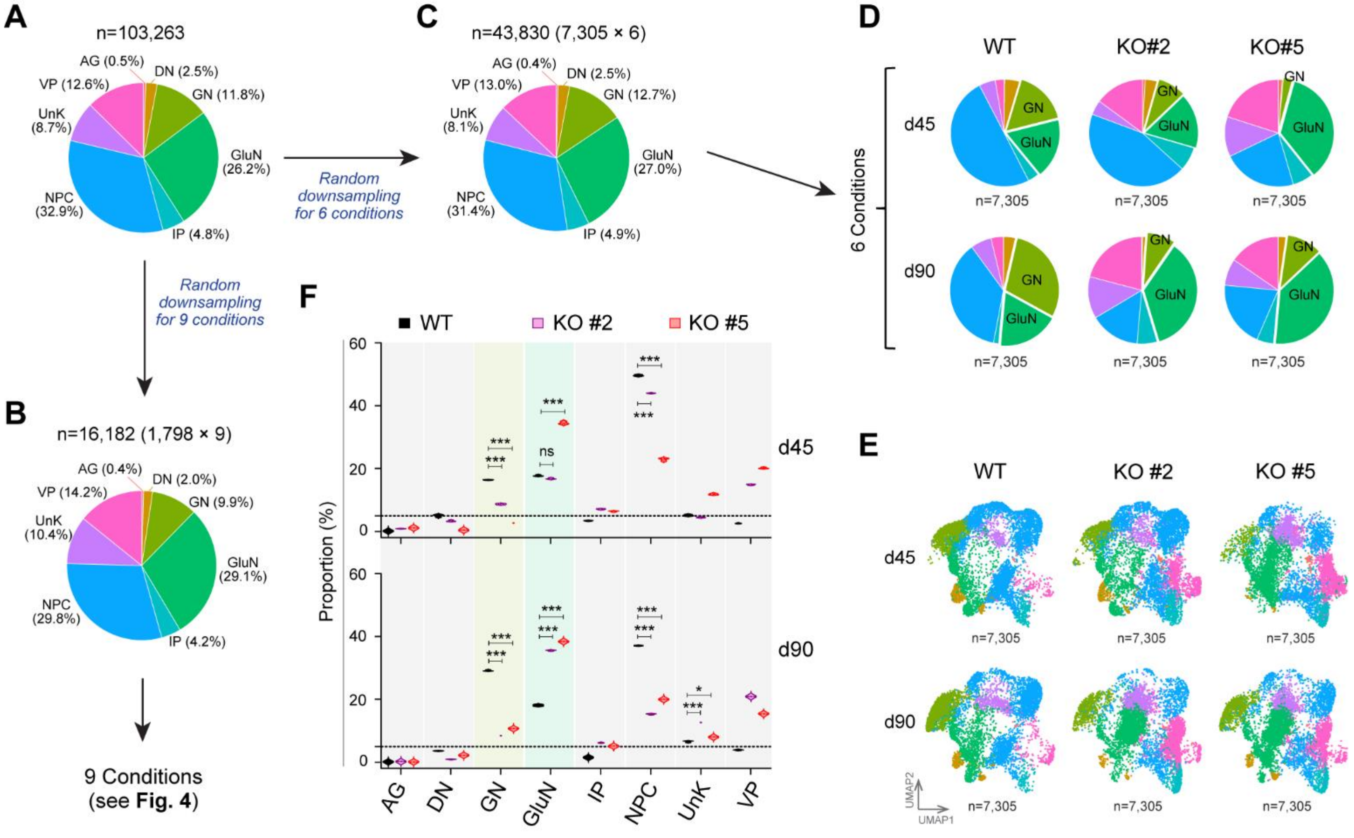

**Figure S9.** Gene ontology (GO) enrichment for (**A**) Cellular Compartment and (**B**) Molecular Function using the 221 differentially expressed genes in UnK cells from the *FNDC4* KO to WT forebrain organoids.

**A**

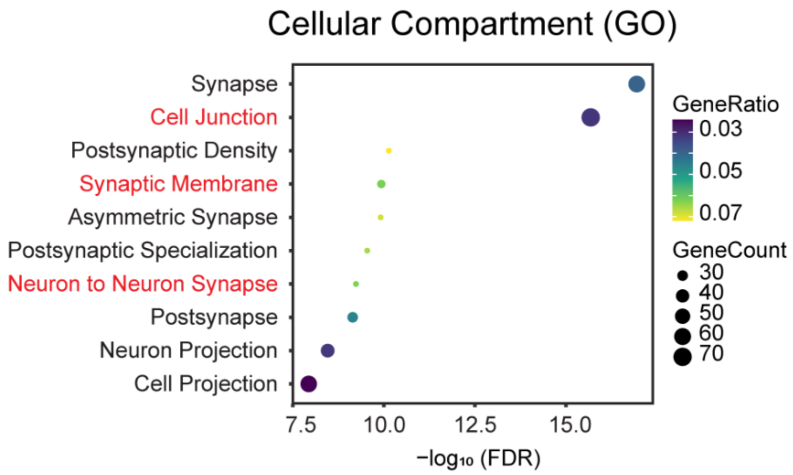


**B**

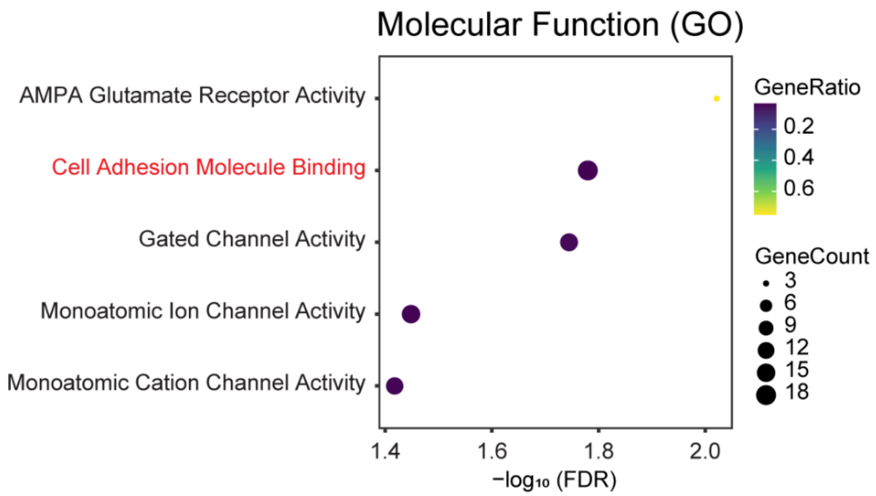
